# Robust software development practices improve citations of RNA-seq tools

**DOI:** 10.1101/2025.09.05.674580

**Authors:** Sanskruti Sharma, Ecem İlgün, Tuna Okçu, Vladyslav Ostash, Vitalina Bashynska, Can Alkan, Elisha Myoung, Naznin Nahar Alpana, Karishma Chhugani, Dhrithi Deshpande, Alex A. T. Bui, Kyle Ellrott, Olga Burduniuc, Tatiana V. Tatarinova, Viorel Munteanu, Paweł P Łabaj, Dumitru Ciorba, Viorel Bostan, Mihai Dimian, Matteo Pellegrini, Mohammed Alser, Serghei Mangul

## Abstract

RNA sequencing (RNA-seq) has emerged as an exemplary technology in biology and clinical applications, offering a crucial complement to other transcriptomic profiling protocols due to its high sensitivity, precision, and accuracy in characterizing transcriptomes. However, the rapid proliferation of RNA-seq tools necessitates the adoption of robust software development practices. Such development underscores the critical need to examine how RNA-seq tools are developed, maintained, and distributed; and whether the data they generate is reproducible as all of these factors are essential for ensuring software reliability, transparency, and trust in scientific findings. We conducted a comprehensive assessment of 434 RNA-seq tools developed between 2008 and 2024, categorizing them based on the type of analysis they perform. Our evaluation encompassed their software development and distribution methodologies, as well as the attributes contributing to their widespread adoption and dependability within the biomedical community, which were quantified by factors such as package manager availability, containerization, multithreading support, documentation quality, and inclusion of example datasets. Our findings establish the first documented positive association between rigorous software development practices and their adoption of published RNA-seq tools as measured by citations (Mann-Whitney U test, p-value = 4.9 × 10^⁻26^). By identifying key characteristics of widely adopted software, our findings guide developing robust and user-friendly RNA-seq tools, thereby reinforcing the call for rigorous community-wide standards.

## Introduction

RNA-sequencing (RNA-seq) is a robust technique that leverages modern sequencing technologies to comprehensively characterize and quantify an organism’s RNA expression, including messenger RNA (mRNA) and total RNA^1,2^. This technique has revolutionized biomedical research by significantly enhancing our ability to analyze the transcriptome, providing unparalleled insights into gene expression and its intricate regulation^3^. Since its inception, RNA-seq has become a leading technology in fundamental biology and clinical applications. It demonstrates superior precision and accuracy compared to more established transcriptomic profiling procedures such as microarrays, which rely on predefined sequences and hybridization for detection^4,5^. As summarized in the US FDA SEQC study, RNA-seq can be used as a versatile tool for relative expression profiling in various applications, provided that sufficient read depth and an appropriate analysis pipeline are chosen^6^.

The continued growth of RNA-seq technologies over the past two decades has led to broad adoption and a rapid increase in the availability of RNA-seq tools. As more tools are developed, implementing robust software development practices becomes increasingly crucial^1,5^. Practices in software development^7^ (e.g., continuous updates with distinctive versioning, and archival stability of tools), robust core functionality (multithreading capacity and user interface) and open science practices (e.g., tool accessibility, easy-to-read user manuals and sample datasets, and easy software installation through packaging and containerization) are necessary to ensure software reliability, transparency, and reproducibility^8,9,10^.

To this end the US FDA SEQC study and accompanying data collection represent a significant milestone in advancing RNA-seq tools. Leveraging datasets of unprecedented size and complexity, along with a diverse set of independent assessment metrics introduced by this study, form a critical resource for benchmarking future RNA-seq developments^11,12^. It has allowed for a comprehensive investigation of 278 representative RNA-seq data analysis pipelines that has resulted in developing guidelines for users to select sensible RNA-seq pipelines for the improved accuracy, precision, and reliability of gene expression estimation and as a follow-up to the improved downstream gene expression-based prediction of disease outcome^13^.

Still, even with established best practices in bioinformatics, several RNA-seq tools fail to adhere to best practices in software development and open science principles. This creates hurdles in installation, compromises performance, and limits reproducibility and accessibility, consequently reducing uptake of such tools by the biomedical community^7,14,15,16,17^. The extent to which scientists incorporate proper software development and open science practices in existing RNA-seq tools remains unknown and has not been previously evaluated.

We aim to rigorously study this issue by conducting a comprehensive evaluation of RNA-seq tools developed between 2008 and 2024, specifically assessing their software development and open science practices. Our primary goal was to understand how these tools were made available to the biomedical community, and assess the relationship between tool adoption, applied software development, and open science practices. Software development practices are evaluated by assessing various features, including the availability of multithreading support, user interface, software updates using versioning, and ease of installation through software packaging and containerization. Open science practices were evaluated by assessing tool accessibility, user guide availability, and archival stability for tools. In addition, the license under which the tools were published was analyzed to determine the accessibility and reusability of the software. Along with effective software development practices, it is equally essential for novel tools to follow best practices in benchmarking. Ideally, a newly developed tool would be compared against state-of-the-art RNA-seq tools^18^, but it is currently unknown whether this is widely followed in the bioinformatics community. Hence, in addition to software development practices, we also examined whether RNA-seq tools follow benchmarking practices.

## Results

### Trends in RNA-seq tool development and community adoption

We performed a comprehensive assessment of software development methodologies, adoption trends, open science compliance, and benchmarking strategies across 434 RNA-seq tools developed between 2008 and 2024 (Figure 1a). We classified the tools into 15 distinct domains based on the type of analysis they perform^19^. These 15 domains span from read alignment to differential expression analysis and transcriptome quantification, encompassing the various steps of RNA-seq analysis workflows. Of the 15 domains, small RNA detection and gene annotation did not experience substantial growth in the number of tools. However, the five domains of read alignment, transcriptome quantification, immune repertoire profiling, differential expression, and transcriptome assembly experienced rapid growth, with an average of 2 to 4 new tools developed per year (Figures 1a, 1b).

**Figure 1:**
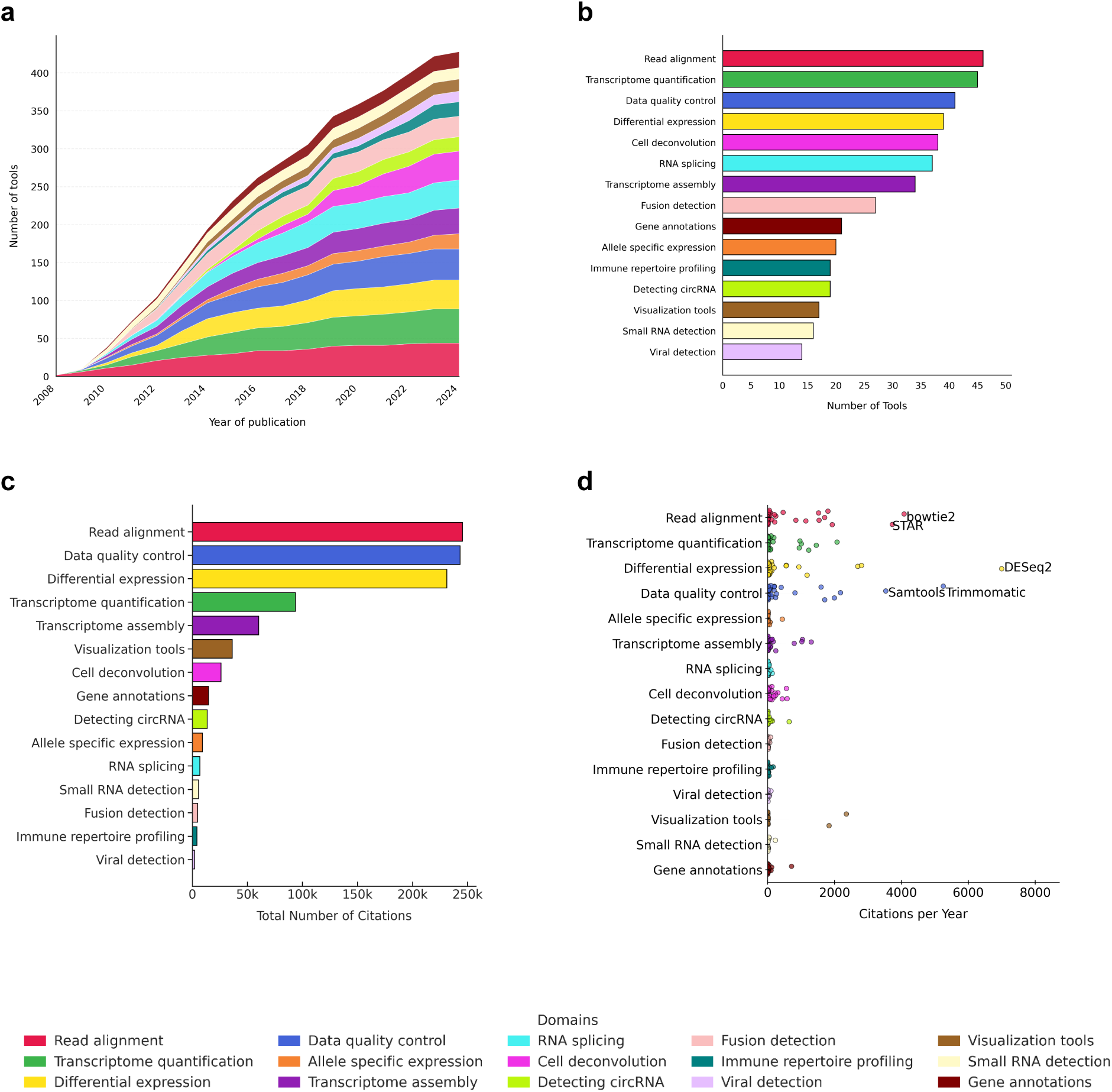
Trends in RNA-seq tool development and community adoption. **a.** Histogram showing the cumulative number of RNA-seq tools developed from 2008 to 2024 (n=434), categorized into 15 domains grouped by the type of RNA seq analysis performed by the tool. **b.** Bar chart showing the number of tools in each domain, with domains ordered in ascending order based on tool count. **c.** Cumulative citation count for all tools within different domains, ordered in ascending order based on citation count. **d.** Citation counts of tools across domains. Tools with the highest citations rates per year are labelled.

To assess the adoption of RNA-seq tools within the biomedical research community, we employed the number of citations per year as a proxy for tool’s adoption. The aggregate number of citations for all tools within a given domain was computed to ascertain the RNA-seq analysis types most frequently used by researchers. Read alignment emerged as the most frequently cited analysis type, with its associated tools collectively accumulating approximately 250,000 citations (Figure 1c). Tools within the data quality control and differential expression domains constituted the second and third most frequently cited types. It is imperative to acknowledge that tools within domains such as read alignment and data quality control possess broader applicability, extending beyond RNA-seq, and encompassing other bioinformatics analyses, including whole-genome sequencing (WGS) and whole-exome sequencing (WES) studies^20^. This wider utility and diversified application likely contributed to their elevated citation counts within the scientific literature.

To identify the most adopted tools, we analyzed the distribution of citations within each domain. Our analysis revealed that in five of the fifteen domains, a single tool accounted for more than 50% of the total annual citations within its respective domain, making each of these tools the most adopted within their field. These tools are GOseq^21^ for allele-specific expression (comprising 60.6% of the domain’s total annual citations), find_circ^22^ for circRNA detection (51.2%), clusterProfiler^23^ 4.0 for visualization (53.7%), miRDeep2^24^ for small RNA detection (53.7%), and Prodigal^25^ for gene annotations (60.6%) (Figure S1a-S1o).

We also observed a high number of citations for several tools from the data quality control and differential expression domains. Within the data quality control domain, tools such as Trimmomatic^26^ and SAMtools^27^ each received over 3,000 citations per year (Figure 1d). Notably, Bowtie2^28^ and STAR^29^, exceeded more than 4,000 citations annually. It is important to acknowledge that while Bowtie2^28^, Trimmomatic^26^ and SAMtools^27^ are versatile bioinformatics tools applicable to various analyses, including RNA-seq, STAR^29^ is specific for RNA-seq. Furthermore, DESeq2^30^ from the differential expression domain stood out, accumulating approximately 7,000 citations per year, making it the most cited tool among all those examined (Figure 1d).

We also evaluated the common benchmarking practices employed by developers of published tools. To analyze this, we examined the original publications in which the tools were presented. Only quantitative, outcome-based benchmarking was considered valid evidence of benchmarking practices; tools that relied solely on qualitative assessments were not included. More than 50% of the RNA-seq tools developed were compared to at least one pre-existing tool at the time of publication (Figure 2a). Surprisingly, benchmarking was not found to be statistically associated with higher citation counts (Figure S2).

**Figure 2:**
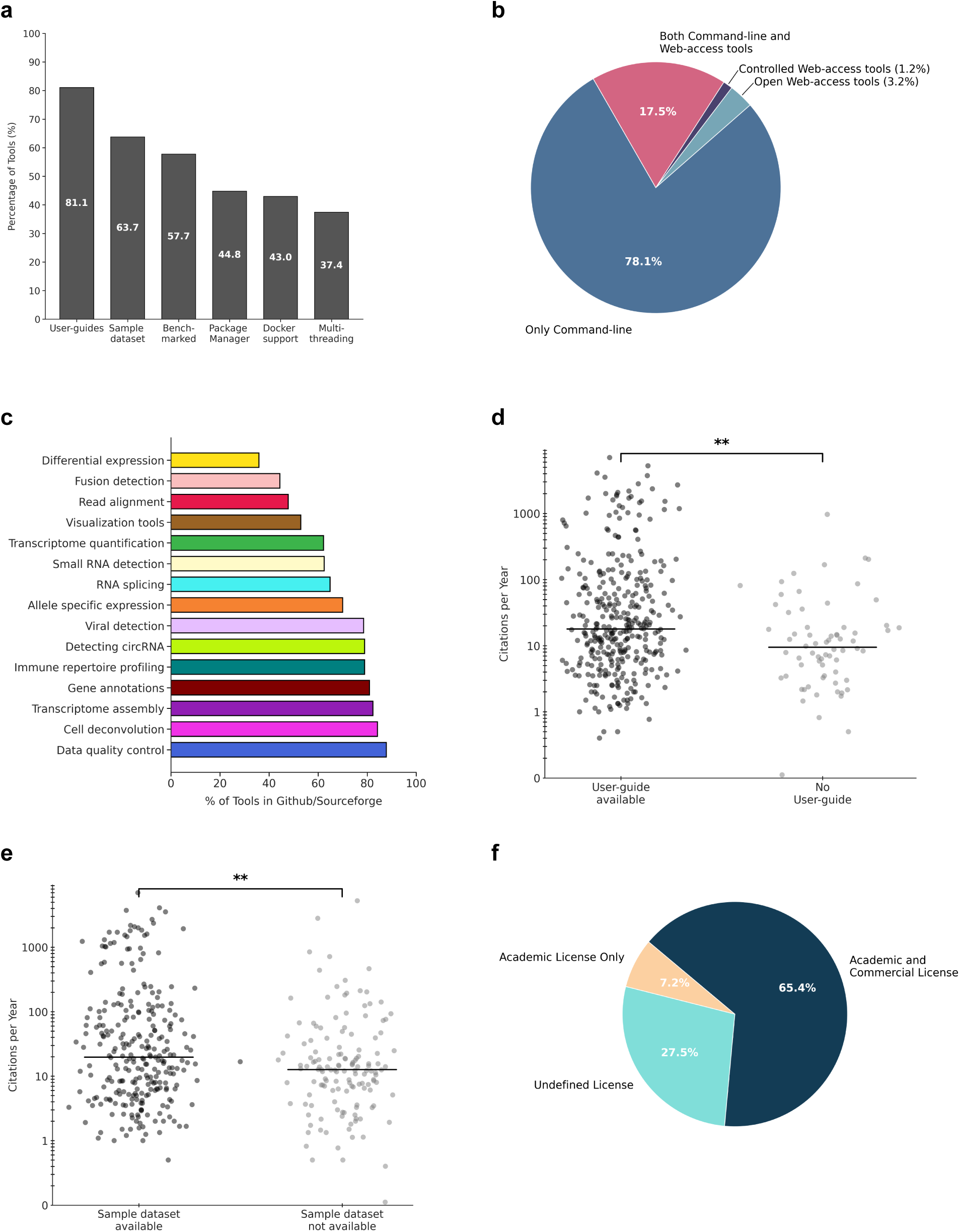
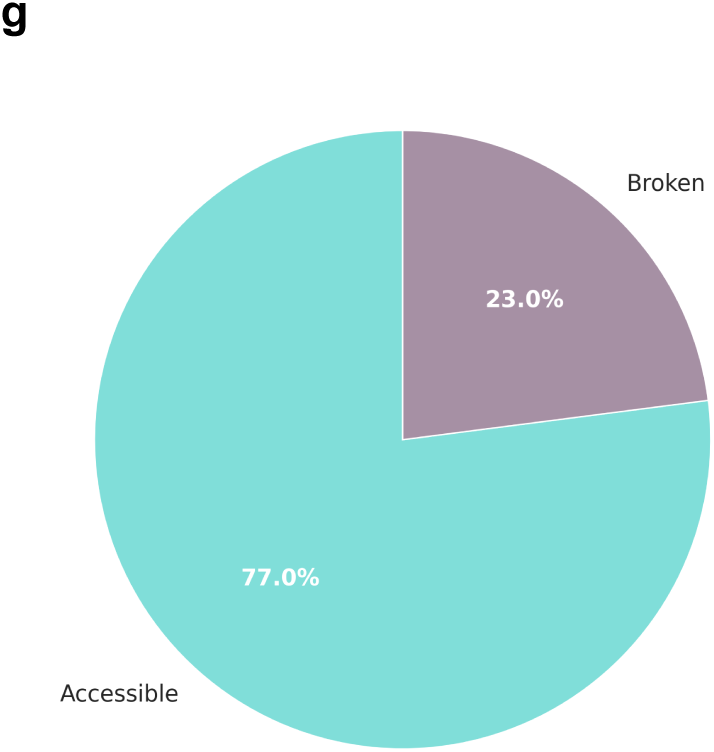
Accessibility and usability of RNA-seq tools. **a.** Bar plot showing the percentage of RNA seq tools that included user guides, availability of sample datasets, adherence to benchmarking standards, support for package management, compatibility with containerization frameworks, and multithreading capabilities during execution. **b.** Pie chart showing distribution of software interfaces among RNA-seq tools, categorized as command-line only, open-access web-based, controlled-access web-based (login required), or hybrid (both command-line and web-based). **c.** Percentage of RNA-seq tools hosted on GitHub or SourceForge, stratified by domain. For each domain, the percentage is calculated relative to the total number of tools in that domain (n = domain-specific tool count). **d.** Tools with user guides are cited significantly more often than those without (Mann - Whitney U test, *p* = 1.10 × 10^⁻3^). **e.** Tools with sample datasets are cited significantly more often than those without (Mann-Whitney U test, *p* = 3.79×10^-3^). **f.** Pie chart showing the percentage distribution of licenses used by developers of RNA seq tools. Tools in the ‘Undefined licenses’ category refer to tools for which no license information was provided or publicly available. **g.** Pie chart showing percentage of RNA-seq tools with accessible vs. broken or links at the time of analysis.

### Accessibility and usability of RNA-seq tools

Enhancing the accessibility and usability of RNA-seq tools is crucial for their widespread adoption within the biomedical community, particularly among researchers with varying levels of computational expertise. Here, we explore the critical features that enhance the usability of RNA-seq tools, examining how elements like the software interface, archival stability, version control, user guides, and sample datasets influence their adoption. Consistent updates, responsive developer support, and the ability to adapt to evolving sequencing technologies are crucial for an RNA-seq tool’s long-term adoption.

To thoroughly assess the usability of RNA-seq tools, we initiated our analysis by characterizing the prevalent interface modalities adopted by researchers, a crucial consideration for developing any novel RNA-seq tool^31^. Our methodology involved a systematic review of each tool’s publicly available source code and its accompanying developer documentation to ascertain its primary interface paradigm: command-line interface (CLI), web-based graphical user interface (GUI), or a hybrid approach. Tools available through Galaxy were also classified as web-based, as Galaxy enables them to be accessed and run via a GUI aligning with the criteria for web-based accessibility. A significantly smaller proportion, specifically 89 tools, provided a dedicated web-based interface. Interestingly, approximately 17% of the tools offered dual functionality, providing both web-based and command-line interfaces, catering to a broader spectrum of user preferences and computational proficiencies (Figure 2b, S3). Furthermore, our investigation into web-based tools highlighted a notable trend regarding accessibility. Roughly half of these web-based tools required user credentials, indicating a controlled access model (Figure S3). This necessitates users to sign up for an account or subscription, potentially limiting open access compared to freely accessible web applications.

Across all 15 domains of RNA-seq analysis, visualization tools emerged as the domain with the highest concentration of web-based tools. This was closely followed by the domains of data quality control and small RNA detection. On the other hand, all tools within the domains of viral detection, gene annotation, circRNA detection, transcriptome assembly, and RNA splicing exclusively provided a command-line interface (Figure S3).

To gain a deeper understanding of developer engagement and version control practices following the publication of a tool, we analyzed its software release frequency. Over 65% of RNA-seq tools used GitHub^32^ and/or SourceForge^33^ not only for sharing the source code but also for sharing compiled binaries of their software and/or releasing updated versions (Figure 2c). The remaining tools were hosted on alternative platforms, such as institutional or laboratory webpages, Bitbucket^34^, or other repositories. In domains such as read alignment, data quality control, and transcriptome quantification, approximately 50% of all tools were hosted on GitHub^32^ and/or SourceForge^33^, indicating a strong reliance on these platforms for source code hosting, version control, and distribution (Figures 2c, S4a and S4b). The domain that contained tools with the highest number of releases per year was differential expression, data quality control, and read alignment, with a mean of approximately 2.96 ± 0.03 releases per year for a given tool in the domain. The small RNA detection and fusion detection domains contained tools with the fewest updates over the years. (Figure S5).

Accessible and well-structured documentation, including practical examples, troubleshooting guides, and tutorials, is vital for empowering users to maximize their tool utilization. In this section, to thoroughly evaluate the usability of RNA-seq tools, we assessed the availability of user documentation, including user guides, manuals, and tutorials that provided structured methodologies for tool operation and comprehensive troubleshooting protocols. Such effective documentation is critical for researchers with diverse computational proficiencies to effectively leverage complex tools and enhance user experience^35^. This is a significant driver of the broader adoption of bioinformatics across the interdisciplinary landscape of life sciences and computational research^36^. Approximately 80% (349 tools) of examined RNA-seq tools provided user guides (Figure 2a), demonstrating a community-wide recognition that comprehensive documentation is essential for ensuring accessibility and effective utilization.

To quantitatively validate this assertion, we conducted a statistical analysis to ascertain the relationship between the availability of comprehensive user guides and a tool’s adoption. Our results demonstrate a statistically significant association, with the presence of user guides being linked to a notable increase in the annual citation count for an RNA-seq tool (Mann-Whitney U test, p-value = 0.001) (Figure 2d). This empirical evidence strongly supports the notion that investing in clear, accessible documentation directly translates into greater scientific visibility within the research community.

Along with user guides, the availability of a sample dataset also provides users with insights into the results and output of the RNA-seq tool and guidance on how to interpret its results. It also increases the reliability and reproducibility of the software, as the test methods utilized by developers in the source code often serve as a sample dataset to validate the core functionality of the tool^10^. We found that more than 60% of RNA-seq tools provide sample datasets in their source code repository (Figure 2a). The availability of a sample dataset was also associated with a higher number of citations per year (Mann-Whitney U test, p-value 0.00379) (Figure 2e).

Next, to ascertain the legal accessibility and alignment with open science of RNA-seq tools, we examined the licensing models under which each tool was released, as indicated in its most updated source code and/or publication. The selection of an appropriate software license is significant, as it directly fosters software innovation by enhancing accessibility for budding researchers and contributing to open science practices^37^. Open licenses facilitate user access and distribution of scientific software, thereby catalyzing opportunities for novel scientific discoveries and technological advancements^38^. Our analysis revealed several key trends in licensing. More than 25% (over 170 tools) lacked explicit or easily identifiable license information (Figure 2f). This absence creates considerable legal ambiguity, potentially hindering researchers’ willingness to adopt, modify, or redistribute these tools due to uncertainties regarding usage rights and intellectual property. This also limits the potential for community contributions and long-term sustainability of software tools^38^. More than 65% (286 tools) of the tools opted for permissive open-source licenses that explicitly permit both academic and commercial utilization (Figure 2f). An example is the GNU General Public License (GPL), a “copyleft” license that ensures users the freedom to run, study, share, and modify software, while requiring any derivative works to remain under the same open license: an approach adopted by around 35% (124 tools) of all RNA-seq tools in our dataset (Figure S6). The Apache License, a highly permissive license, allows free use, modification, and distribution for any purpose, including commercial, with minimal restrictions. The Artistic License, primarily used for Perl projects, balances freedom with some control over derivative works^39–43^. The remaining tools were published under licenses that strictly limit their use to academic or non-commercial purposes (Figure 2f). While these licenses can protect intellectual property in non-commercial settings, they inherently restrict broader adoption, particularly for industrial or commercial purposes.

Lastly, we evaluated the archival stability of the uniform resource locators (URLs) provided by scientists when publishing novel tools, examining how frequently these tools are updated on their hosting platforms and the number of versions available. This assessment aimed to determine whether developers ensure continued access or if the links become inaccessible or nonfunctional over time. Continued accessibility to a tool through stable software links is a fundamental first step toward developing a computational resource that ensures sustained reach within the community^10^. We found that out of 434 tools, more than 75% offered accessible links (Figures 2a, 2g). However, this indicated that software links were broken for over 100 tools out of 434, making the tool inaccessible. For such tools, we tried to find alternative links to extract data. As expected, the presence of an accessible software link was associated with higher citation counts (Mann-Whitney U test, p-value 0.0186) (Figure S7).

### Robust software development practices improve citations of RNA-seq tools

Software development practices that prompt long-term usability, transparency and reproducibility are widely regarded as robust and reliable in supporting high-quality credible research outcomes^9,10^. Robust software development practices are increasingly recognized as pivotal for enhancing the adoption and citation count of RNA-seq tools within the dynamic biomedical research landscape. To ensure software quality, reliability, and reproducibility which are the foundational tenets of credible scientific research, it is imperative that newly developed RNA-seq tools rigorously adhere to established best practices in scientific software engineering^9^. We investigated how software development characteristics, such as support for multithreading capabilities, package managers, and containerization, enhance a tool’s computational performance and contribute to its increased utilization within the scientific community, ultimately leading to greater academic recognition. By comprehensively evaluating how these practices are implemented across RNA-seq tools, we aim to provide empirical evidence for their collective adoption on a tool’s citation rate, perceived utility, and overall contribution to the biomedical research ecosystem, offering critical insights for both developers striving to create well-engineered tools and researchers seeking reliable and sustainable solutions for their RNA-seq analyses.

As a critical first step in evaluating the practical usability and reproducibility of RNA-seq tools, we systematically assessed the availability of tools within established package managers. Package managers are indispensable for automating the tedious processes of software tool installation, configuration, and ongoing maintenance. They significantly streamline the setup process and expertly resolve complex inter-tool dependencies. They play a defining role in ensuring the consistency and accessibility of computational environments for RNA-seq analysis^44^.

Despite the profound advantages offered by package managers, our comprehensive analysis revealed a notable gap: more than 55% (238 tools) of the surveyed RNA-seq tools were not packaged (Figures 2a, S8). This necessitates manual installation and dependency resolution, introducing considerable friction and potential for error, particularly for researchers who may not possess advanced computational skills^7,14^.

Analysis of tool distribution across RNA-seq domains revealed distinct patterns. The differential expression domain exhibited the highest proportion of tools deployed via package managers, indicating a recognized advantage among developers in this area for streamlined software deployment. This was closely followed by domains such as cell deconvolution and gene annotations, suggesting an evolving trend towards improved packaging in core analytical areas. Conversely, in the viral detection domain, over 80% of tools lacked support for a package manager (Figure 3a). Some tools could be deployed via multiple package managers. For this we counted each instance of package manager separately. Among all the tools that implemented package managers, Anaconda^45^ was the most widely used, supporting the distribution of over 95 RNA-seq tools (>50%) (Figures 3a, 3b and S9). Data quality control emerged as the domain with the highest number of Anaconda-packaged tools, with an average of approximately nine tools per domain utilizing Anaconda^45^ for distribution. On the other hand, the visualization domain has the least number of tools that were packaged using Anaconda^45^ (Figure 3a). The availability of package managers was also found to be associated with a significant increase in the number of citations (Mann-Whitney U-test, p-value 1.04 × 10^-21^) (Figure 3c).

**Figure 3:**
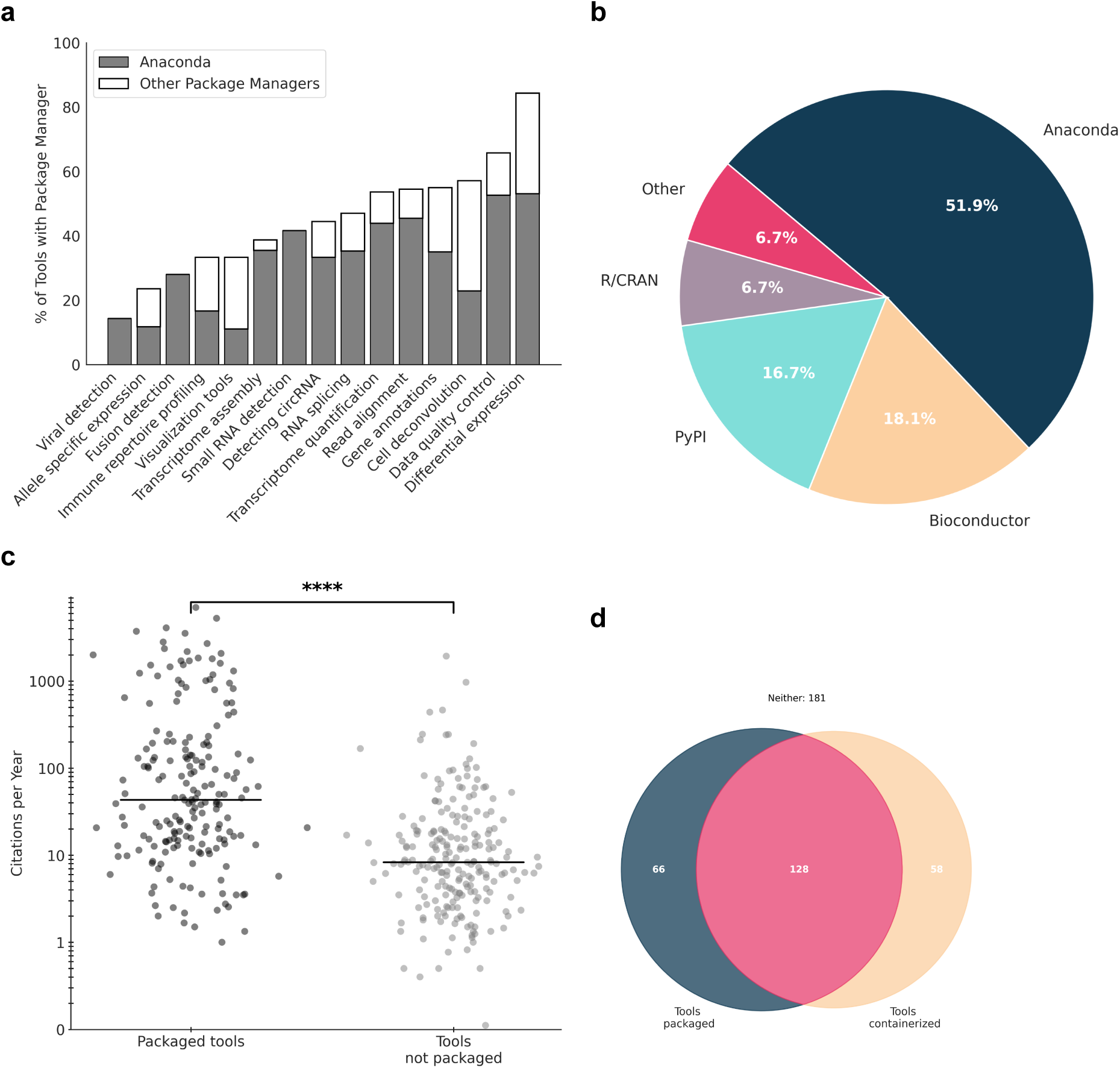
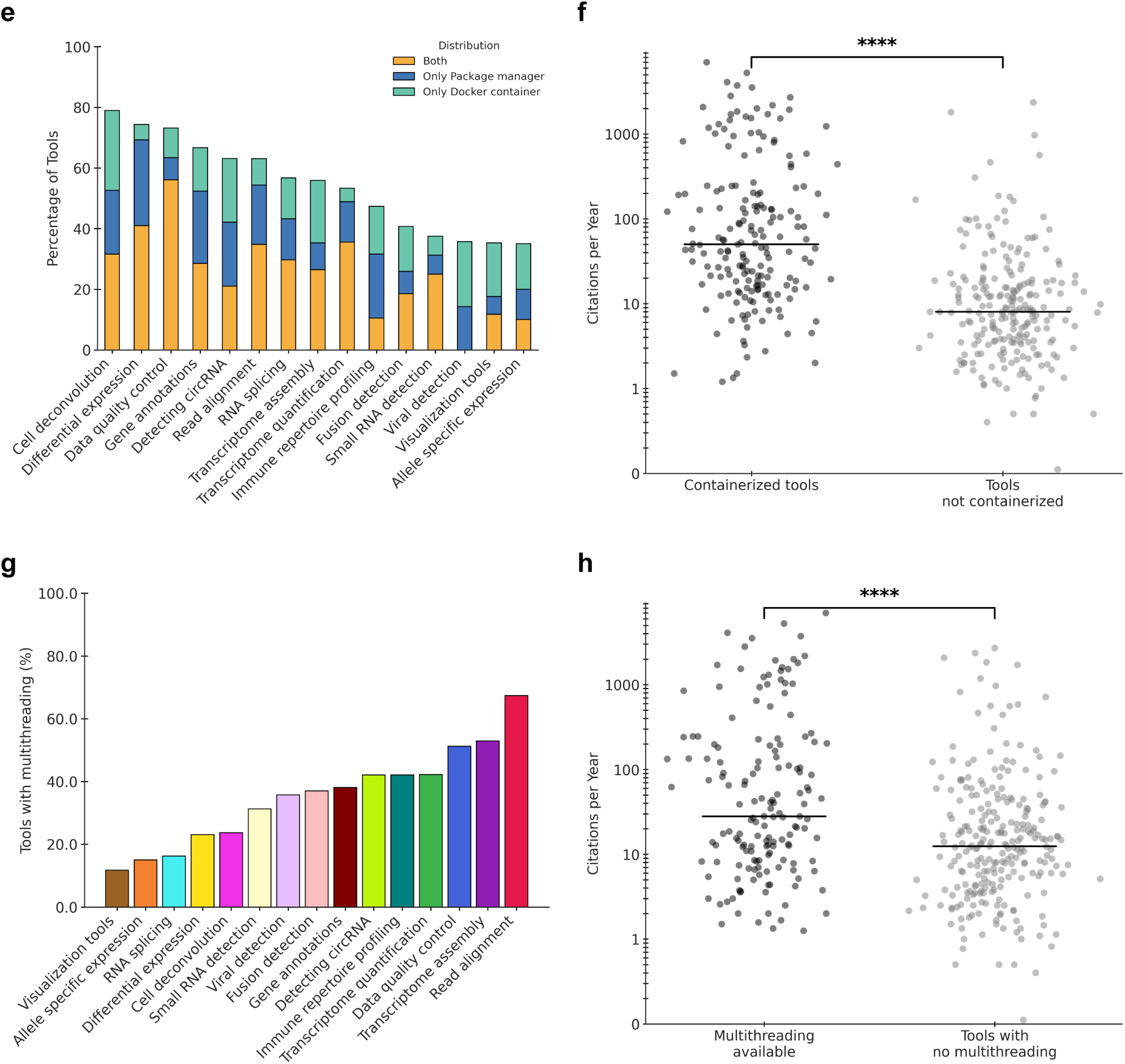
Software development practices of RNA-seq tools. **a.** Stacked bar plot showing the percentage of RNA-seq tools packaged, with those installable via Anaconda shaded in grey. **b.** Pie chart showing distribution of different package management systems across all command-line RNA-seq tools. **c.** Tools with package manager support are cited significantly more often than those without (Mann - Whitney U test, p = 1.04 × 10^-21^). **d.** Venn diagram showing overlap of tools with both package manager support and Docker containerization, and those with neither. **e.** Percentage of tools in each domain classified by their support for Docker, package managers, both, or neither. **f.** Tools with Docker containerization are cited significantly more often than those without (Mann-Whitney U test, p = 4.96 × 10^⁻26^) **g.** Bar plot showing percentage of tools with multithreading capacity during execution (n = domain-specific tool count) **h.** Tools with multithreading capability are cited significantly more often than those without (Mann-Whitney U test, p = 1.03 × 10^-6^).

Building upon our assessment of package manager support, our next step focused on examining the prevalence of containerization via Docker^46^. This advanced containerization technology further enhances reproducibility and portability by allowing applications to run independently from the host operating system, thereby boosting software speed and efficiency and facilitating faster, more reliable deployments. While package managers simplify installation, Docker^46^ containers ensure consistent execution across diverse computing environments, addressing a crucial layer of computational reproducibility^44,47^. Our analysis revealed a significant adoption of Docker containerization within the RNA-seq ecosystem. More than 40% (186 tools) of all RNA-seq tools had Docker^46^ containerization capacity, indicating a growing recognition among developers of the benefits of encapsulated environments for complex bioinformatics workflows. Consistent with the trends observed for package manager support, the data quality control domain once again emerged at the forefront, with the highest number of tools that could be executed within a Docker^46^ container with approximately 70% (30 out of 43 tools in this domain) providing this capability. This strong adoption of data quality control reinforces the domain’s commitment to robust and user-friendly deployment mechanisms. However, a notable disparity became evident. Less than 25% (14) of the tools in viral detection and allele-specific expression domains had Docker^46^ containerization capacity. (Figure S10). Interestingly, we observed that more than 65% (125) of the tools with a package manager available also had Docker^46^ containerization capacity (Figure 3d). The data quality control domain exhibited the highest number of packaged and executable tools, all of which were deployed via a Docker^46^ container. By contrast, viral detection tools lacked support for Docker^46^ containers and package managers (Figure 3e). Many older tools predate the introduction of package managers such as Anaconda^45^ and container platforms like Docker^46^, and thus could not incorporate these technologies.

Similarly, we conducted a statistical analysis to ascertain the relationship between Docker^46^ containerization capacity and its influence on citation count. We found a statistically significant association, with the tools that could be executed within a Docker^46^ container being associated with a significant increase in the citation count and tool adoption (Mann-Whitney U test, p-value = 4.96 × 10^-26^) (Figure 3f).

We proceeded to investigate the internal computational efficiencies of RNA-seq tools, with a particular focus on implementing parallelization methods, specifically multithreading. Multithreading is a critical design choice that enables software to leverage multiple processing cores concurrently, thereby increasing computational efficiency and facilitating faster analysis of large-scale genomic datasets. This capacity for speed is paramount in a field where data volumes continue to rapidly expand^48^.

Our findings indicate that multithreading has been widely adopted across diverse categories of RNA-seq tools, reflecting a broad understanding among developers of its importance for performance. Over 36% (157 tools) of all RNA-seq tools surveyed offered multithreading support. The read alignment domain, which often involves computationally intensive tasks of mapping millions or billions of short reads to a reference genome, exhibited the highest proportion of tools enabling multithreading, a testament to the necessity of parallel processing in this foundational step. Following the read alignment domain, approximately 50% of tools in domains such as transcriptome assembly (17 out of 33 tools) and data quality control (19 out of 41 tools) also incorporate multithreading capabilities (Figure 3g). We observed that multithreading capacity was significantly associated with higher citation counts for RNA-seq tools (Mann-Whitney U test, p-value = 1.06 × 10^-6^) (Figure 3h).

### Best practices for RNA-seq software development

Here we summarize the best RNA-seq software development practices that enhance reliability, usability, transparency, and accessibility (Box 1). Software reliability is closely linked to implementing robust computational features and their efficiency, such as package manager integration, support for multithreading, containerization with Docker^46^, and using a command-line interface to facilitate deployment in high-performance clusters. Developers should incorporate these features to enhance the stability and scalability of the software, as well as to ensure ease of installation, compatibility across environments, and efficient resource utilization^49^. Statistically significant findings in this study also support the claim that Docker containerization and the multithreading capacity of a tool directly translate into higher adoption rates of the tool (Figures 3f, 3h). This finding indicates that tools engineered for computational efficiency and speed are more likely to be adopted by users and integrated into research pipelines, ultimately contributing to broader scientific adoption.

#### Box1: Recommended Best Practices for RNA-seq Software Development

Based on our comprehensive analysis, we delineate critical best practices for RNA-seq software development that are pivotal for enhancing tool reliability, usability, transparency, and accessibility. Adherence to these guidelines can significantly bolster the bioinformatics community’s capacity for reproducible and efficient research.

1. Software Reliability and Performance: Robust computational features and efficiency are foundational to reliable software.

- Performance Enhancement Techniques: To maximize performance, developers should implement advanced techniques including multithreading support, and leveraging Graphics Processing Units (GPUs). These significantly improve execution speed and resource utilization.
- Containerization: The use of containerization technologies (e.g., Docker, Singularity) is essential. Containers encapsulate the application and its dependencies, ensuring consistent performance, boosting speed and efficiency, and facilitating cross-environment compatibility by eliminating issues related to disparate operating systems and software setups.
- Command-Line Interface (CLI): The provision of a robust CLI is crucial for deployment in high-performance computing (HPC) environments, enabling automation and scripting for large-scale analyses. Where appropriate, a user-friendly graphical user interface (GUI), such as a web-based interface, can also be offered to enhance user interaction.
2. Usability and Accessibility: Clear and comprehensive documentation, alongside streamlined installation, is paramount for widespread adoption.

- Comprehensive Documentation: A well-structured README file in the source code repository should provide detailed instructions for downloading, installing, and running the tool. This must be complemented by a detailed user guide covering alternative options, parameter values, and troubleshooting tips, catering to users with varying computational expertise.
- Illustrative Examples: Developers should include a sample dataset to aid user understanding and verify functionality. This allows users to quickly assess expected output and confirm the tool’s completeness and correct operation.
- Package Manager Integration: To streamline the user experience, tools should be packaged using widely adopted package managers (e.g., Anaconda, Bioconductor, PyPI, CRAN). This automates installation, resolves complex dependencies, and simplifies maintenance, making the tool significantly more user-friendly.
3. Transparency and Open Science: Public availability and clear licensing foster collaboration and trust.

- Public Code Release: Once the source code is thoroughly tested and ready for publication, developers of RNA-seq tools should publicly release it on well-maintained platforms such as GitHub. This not only fosters open science but also ensures the code is accessible for review, contribution, and long-term preservation.
- Software Licensing: The repository must explicitly specify a software license (e.g., GPL, MIT). This defines terms of use, encourages broader adoption, and supports innovation within the community.
4. Sustained Engagement and Validation: Ongoing maintenance and empirical validation are critical for long-term viability and credibility.

- Active Maintenance and Versioning: Beyond the initial release, continuous engagement is vital. Developers should actively maintain their software tools, providing clear descriptions of differences between versions. This ongoing development and transparent versioning build user trust and ensure the tool’s long-term viability.
- Benchmarking and Performance Validation: To demonstrate value and validate performance, developers should rigorously benchmark their creations against existing state-of-the-art tools. This comparative analysis provides crucial evidence of efficacy, helping users make informed decisions and contributing significantly to the tool’s credibility and adoption within the scientific community.

Usability and accessibility of a software, however, are dictated by a variety of factors. To ensure clear documentation in the source code repository, developers should include a well-structured README file with instructions for downloading, installing, and running the tool. This can be significantly enhanced by a detailed user guide that covers alternative options, parameter values, and troubleshooting tips, catering to users with varying computational expertise. To further aid understanding and verify functionality, developers should include a sample dataset, allowing users to quickly examine expected output^10,50^ and confirm the tool’s completeness^51,52^.

Transparency is essential in software development. Once the source code is well-tested and ready for publication, developers should publicly release it on well-maintained platforms such as GitHub^32^. This practice not only promotes transparency and open science, but also ensures that the code is accessible for review and community contributions. Additionally, the repository should clearly specify a software license (e.g., GPL, MIT) to define the terms of use and encourage broader adoption and innovation^39,40^.

To streamline the user experience, we recommend developers package their tools using widely adopted package managers such as Anaconda^45^, Bioconductor^53^, PyPI^54^, or CRAN^55^. This automates installation, resolves complex dependencies, and simplifies maintenance, making the tool much more user-friendly. In today’s diverse computational landscape, containerization techniques (e.g., Docker^46^, Singularity^56^) are essential^57^. They allow the application to run independently of the host operating system, boosting speed and efficiency while ensuring consistent performance across different environments. This is particularly important for avoiding issues with disparate operating systems and software setups^47^.

Beyond initial release, sustained engagement is key. Developers should actively maintain their software tools and provide clear descriptions of differences between versions. This continuous development and transparent versioning contribute significantly to user trust and the tool’s long-term viability^39^. Finally, to demonstrate the tool’s value and validate its performance, developers should benchmark their method against existing state-of-the-art tools. This comparative analysis provides crucial evidence of efficacy and helps users make informed decisions, contributing to the tool’s credibility and adoption within the scientific community^18^.

## Discussion

We evaluated the software development and open science practices of 434 RNA-seq tools developed between 2008 and 2024. Our investigation highlights an ongoing challenge: while the number of RNA-seq tools has grown substantially over the past two decades, there remains an opportunity for the bioinformatics community to establish and adopt more widely formalized and consistent guidelines to enhance tool accessibility, usability, and reliability.

Our assessment reveals a gap in the software development practices of many RNA-seq tools. For instance, a substantial proportion still lack essential features, such as package manager availability and containerization. Specifically, we found that over 50% of the tools surveyed do not offer package manager support, and a similar proportion lack Docker^46^ containerization. We also found that certain domains, such as visualization tools and small RNA detection, had a higher number of web-based tools compared to other domains. This distribution suggests that developers recognize the benefits of more intuitive, visually-driven interfaces for specific stages of RNA-seq analysis, which often involve initial data inspection and core processing, thereby enhancing access for a wider research audience. Furthermore, over 60% of the tools lack multithreading capabilities. These deficiencies are critical because our study provides compelling evidence that effective software development practices are strongly associated with higher citation counts, which demonstrates greater adoption within the biomedical community (Mann-Whitney U test, p-value 4.9 × 10^−26^). This highlights an urgent need: we must establish and disseminate clear software development guidelines to enhance tool efficiency and help researchers perform reliable RNA-seq analysis. Our findings support previous observations about the challenges in bioinformatics software development^7,14^. We provide concrete evidence of how robust practices influence the adoption of tools. Adhering to established best practices and community guidelines can significantly strengthen the bioinformatics community by enhancing the usability, credibility, and widespread adoption of new tools.

While our study offers a broad perspective, it is essential to acknowledge certain limitations. First, while extensive, our dataset of RNA-seq tools is not exhaustive due to the continuous emergence of novel tools as technology advances. To mitigate this, we systematically categorized state-of-the-art RNA-seq tools. We grouped them by domains, allowing us to identify trends among scientists addressing specific research problems, such as differential gene expression or sequence quality assessment. Second, it is crucial to note that several tools frequently embedded in standard RNA-seq pipelines were not originally developed exclusively for RNA-seq analysis. For instance, alignment tools like Bowtie2^28^ and BWA^58^, initially designed for DNA sequencing, are now widely used for mapping RNA-seq reads. These tools, despite their broader origins, are integral to RNA-seq workflows, and their inclusion in our analysis provides a realistic representation of the tools employed by the community. Another limitation of our study is that we focus solely on the presence or absence of various features or parameters associated with higher citation counts, rather than assessing their quality. For example, we record whether documentation is provided in the source code, but do not evaluate its depth or clarity. Similarly, our study did not incorporate a temporal evaluation of parameters; for instance, we could have examined how the citation frequency of a given tool has varied over time to capture changes in its scholarly impact. While our current approach enables broader coverage, incorporating qualitative analysis in future work could offer additional insights into how these features influence impact. Finally, we recognize that citation count, while a strong indicator, is not the sole determinant of a tool’s adoption. Other factors, such as author-level metrics (e.g., h-index, g-index), journal-level metrics^59,60^ (e.g., impact factor, journal rank), as well as broader community recognition and the strength of a tool’s promotion can also influence its visibility and perceived importance^61^. While our study focuses specifically on software development practices, we acknowledge these broader academic factors. Along with addressing these limitations, future work should also consider all the FAIR principles (Findable, Accessible, Interoperable, Reusable), which extend beyond data to include RNA-seq tools, thereby promoting transparency and reproducibility across all aspects of the research lifecycle.

Our study directly examines how robust software development practices influence citation count. Previous research has explored various factors influencing citations in bioinformatics; for example, Glänzel et al. (2009)^59^ examined journal-level metrics and collaboration patterns, and Perez-Iratxeta et al. (2007)^60^ investigated how shifts in research focus affect citations. More recently, Song M et al. (2014)^61^ considered geographical factors and author-level metrics, and Ibáñez et al. (2009)^62^ utilized machine learning to predict citation count based on journal, author, publication timing, and abstract keywords. While these studies offer valuable insights, and despite the potential for inadvertent errors due to our own manual data extraction, our work delves deeper than prior analyses by examining concrete features of the tools themselves, such as the availability of package managers, Docker^46^ support, and multithreading capabilities. We demonstrate their positive effect on increased adoption and robust software development, providing actionable insights for developers aiming to maximize the reach and adoption of their bioinformatics tools.

We strongly encourage computational biology and bioinformatics researchers to adopt these robust software development practices. Doing so significantly increases the citation count and visibility of their published tools while also meaningfully advancing research quality and promoting best practices in scientific software development across the entire biomedical community. An interesting future direction would be to develop a machine learning model capable of predicting the citation count per year for RNA-seq tools. Such a model could be trained on the key factors identified in this study, including package manager availability, Docker^46^ containerization, year of publication, multithreading support, user guide accessibility, and the presence of sample dataset to quantitatively assess their adoption by the biomedical community.

## Methods

### Protocol for evaluating the design and capabilities of RNA-seq tools

All information regarding the features and characteristics of RNA-seq tools was extracted manually. Our inclusion criteria prioritized tools that explicitly mentioned “RNA” or “RNA-seq” in their title, abstract, or publication. Additionally, we included general-purpose tools that utilized RNA-seq data as input. The primary sources for collecting data about RNA sequencing tools are databases and search engines. Databases such as PubMed and Oxford Academic, and search engines like Google Scholar, were utilized for identifying RNA-seq tools. The RNA-seq tools selected in our analysis fell into two categories: those explicitly developed for RNA-seq data processing and general-purpose bioinformatics tools applicable to bulk RNA-seq analysis. The tools were selected based on an inclusion criterion: they must be able to analyze RNA-seq data and be openly accessible. For each tool, the tool name, year of publication, software interface utilized, package manager, Docker^46^ containerization, multithreading capacity, user guides, sample dataset, link maintenance, and benchmarking practices conducted were studied. For information regarding the software interface, it was noted whether the tools were web-based - with or without controlled access - used a command-line interface, or offered both options. Tools available through Galaxy^63^ were also classified as web-based tools, as it enables them to be run locally via a graphical user interface (GUI), aligning with the criteria for web-based accessibility.

For package managers, information regarding which package manager (Anaconda^45^, Bioconductor^53^, CRAN^55^, PyPI^54^, or other lesser-known package managers, such as Homebrew^64^) was utilized, as well as the number of releases and/or updates that the tool had in each package manager, was also noted. Along with this, citations and citations per year for each tool were also cataloged.

After data extraction, we categorized the tools into 15 domains: allele-specific expression, cell deconvolution, data quality control, detecting circRNA, differential expression, fusion detection, gene annotations, immune repertoire profiling, RNA splicing, read alignment, small RNA detection, transcriptome quantification, transcriptome assembly, viral detection, and visualization, based on the primary function for which the tool was developed.

We curated data from tools published between 2008 and 2024. Data, including the availability of package managers and Docker support, was extracted from publicly available repositories manually, by searching for the tool in the repository and counting the versions released for that tool. We investigated whether the tool was benchmarked against other tools at the time of publication and assessed whether the benchmarking practices conducted were solely qualitative. Various Python libraries, namely Numpy^65^, Seaborn^66^, and Matplotlib^67^, were utilized for data visualization.

### Statistical Analysis

Statistical analysis was conducted to investigate the relationship between various software development characteristics and citation counts. Key attributes such as package manager availability, Docker^46^ containerization, multithreading support, manual and sample dataset availability, user-guide availability, and benchmarking practices were tested and for each feature, citations per year were also evaluated to reveal associations. As the citation count did not follow normal distribution, we used statistical tests that do not assume normality for all comparisons. Specifically, the Mann–Whitney U test was applied to compare citation metrics between groups defined by the presence or absence of each feature, with statistical significance evaluated at a threshold of 0.05. All analyses were performed using Python and relevant scientific libraries (NumPy^65^, Matplotlib^67^, Seaborn^66^, and SciPy^68^). Full statistical results, including test statistics and p-values for each comparison, are available at https://github.com/Mangul-Lab-USC/RNA-seq

## Acknowledgements

We thank the authors of the tools surveyed in this work for providing helpful feedback and verifying the information related to their tool.

## Code availability

All code used in this study is publicly available at: https://github.com/Mangul-Lab-USC/RNA-seq

## Data availability

All data generated and analyzed during this study are publicly available at: https://github.com/Mangul-Lab-USC/RNA-seq

