## Supplementary figures and images for "Robust software development practices improve citations of RNA-seq tools"

### Figure S1a

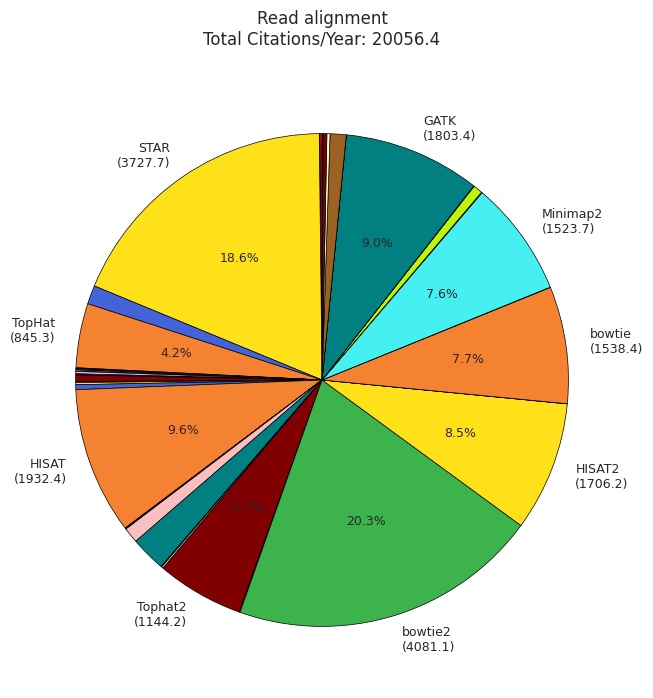

### Figure S1b

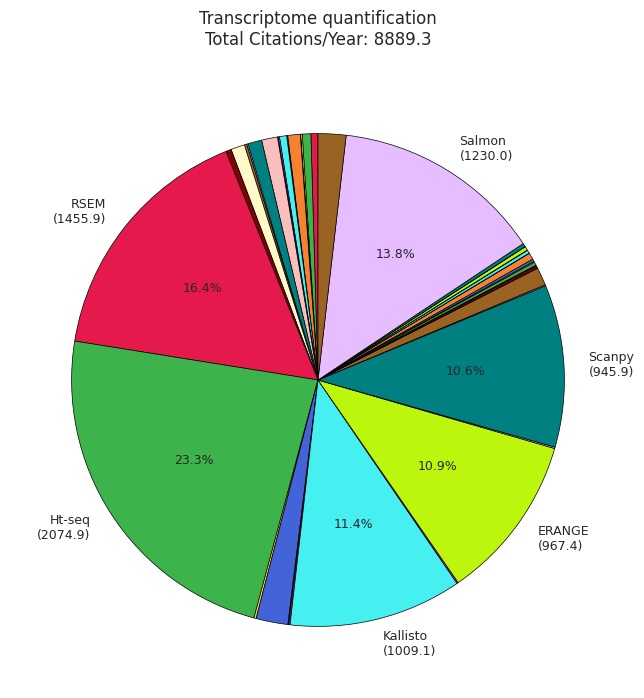

### Figure S1c

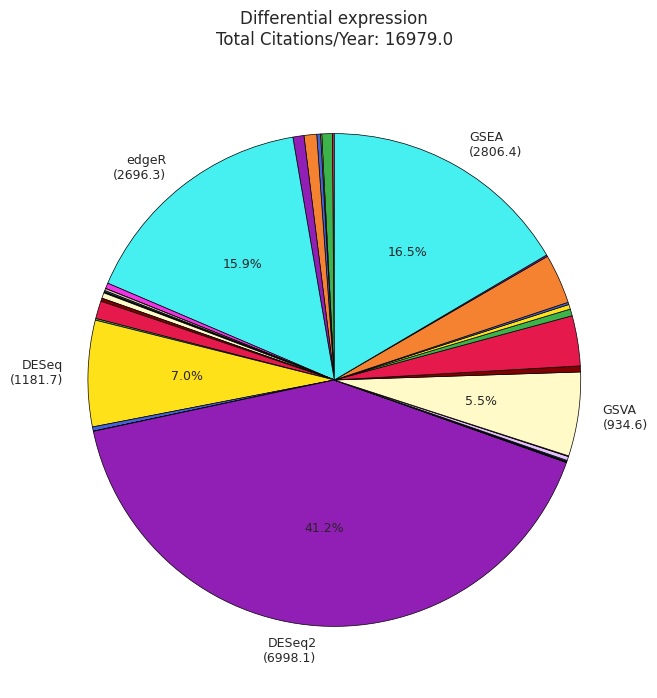

### Figure S1d

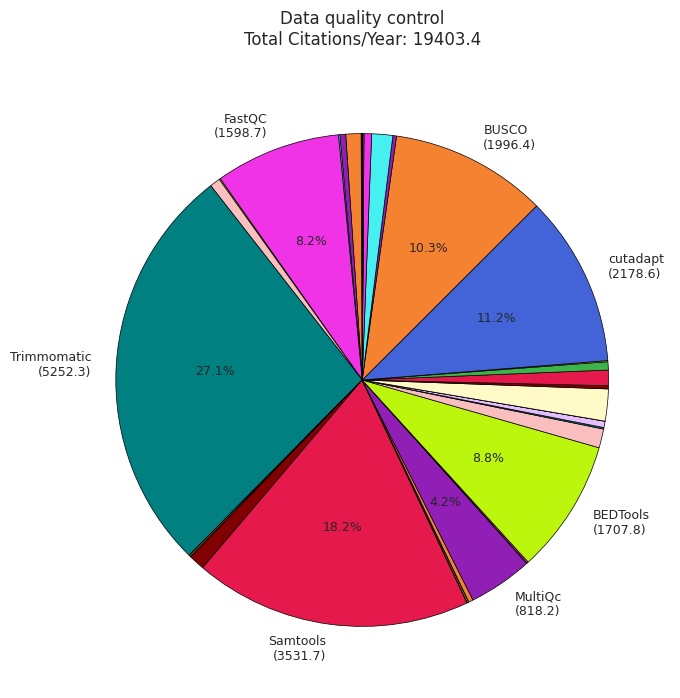

### Figure S1e

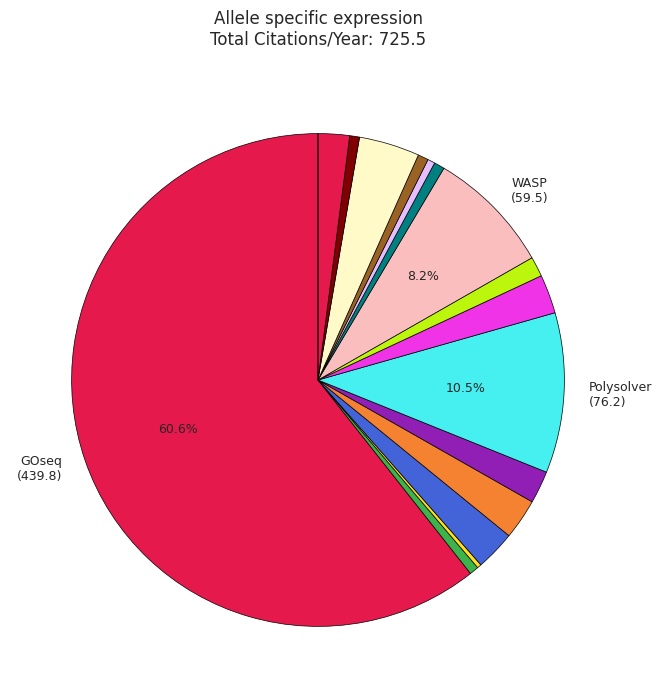

### Figure S1f

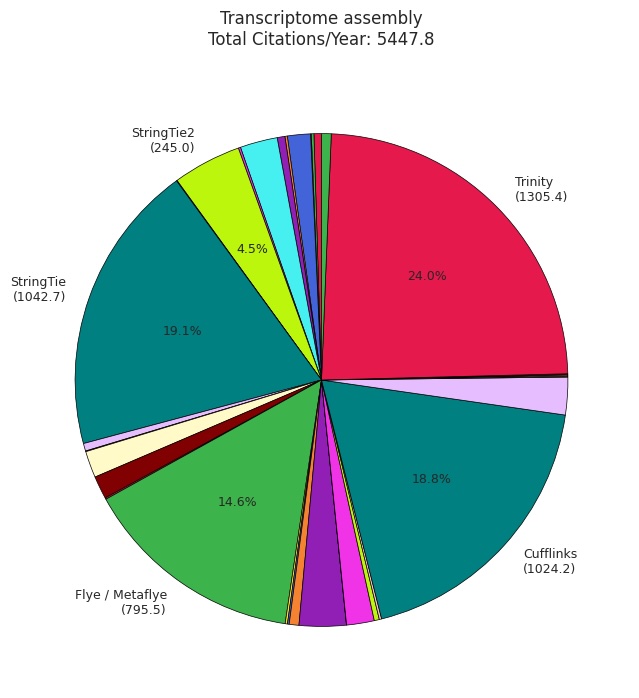

### Figure S1g

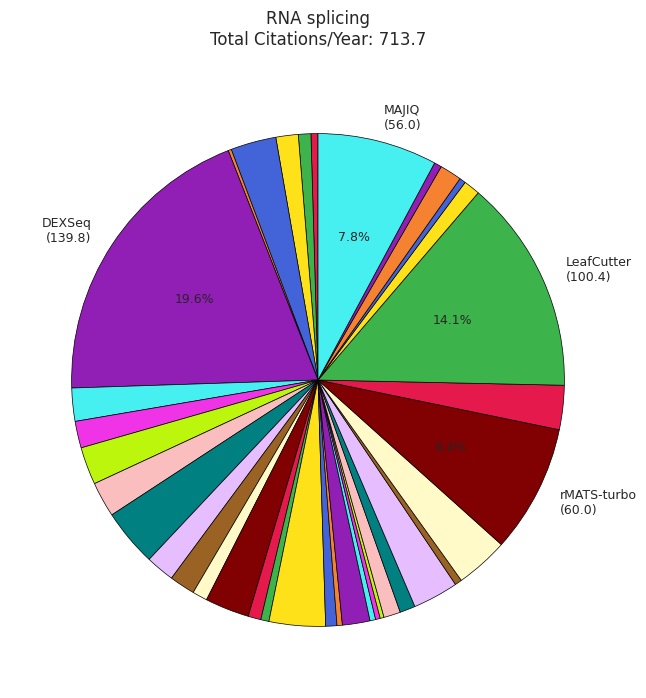

### Figure S1h

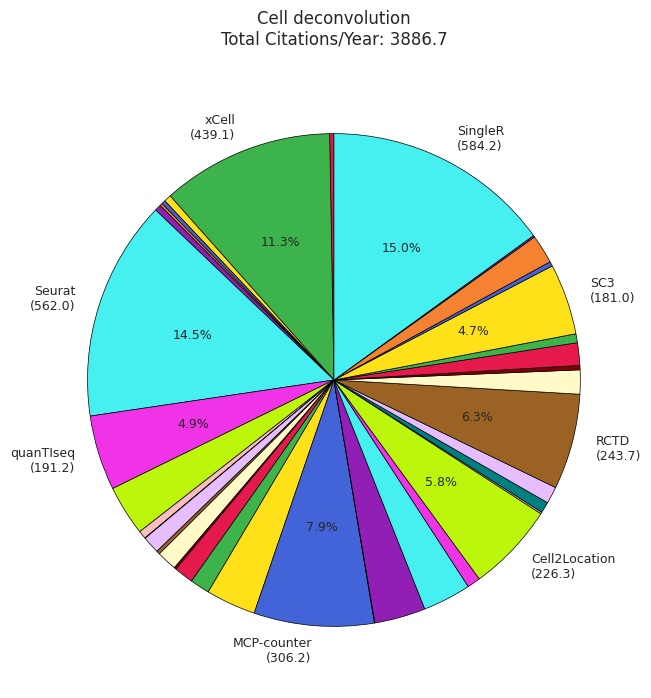

### Figure S1i

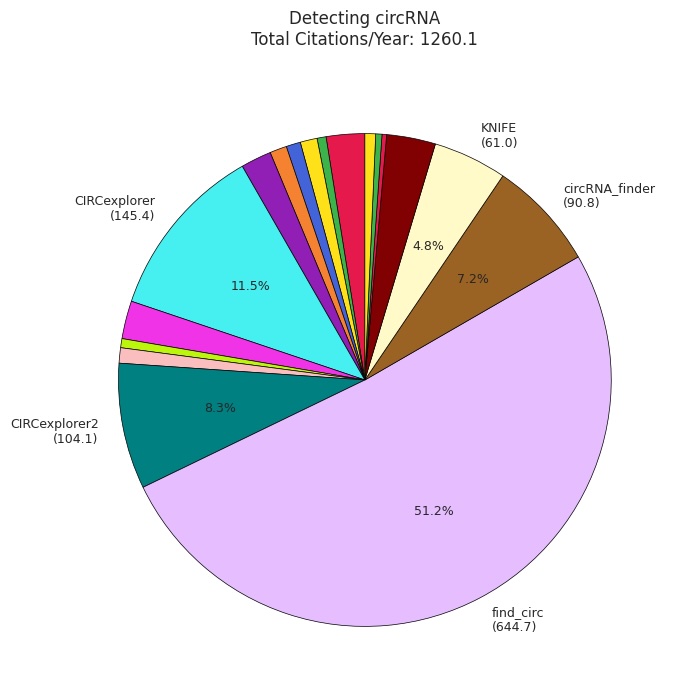

### Figure S1j

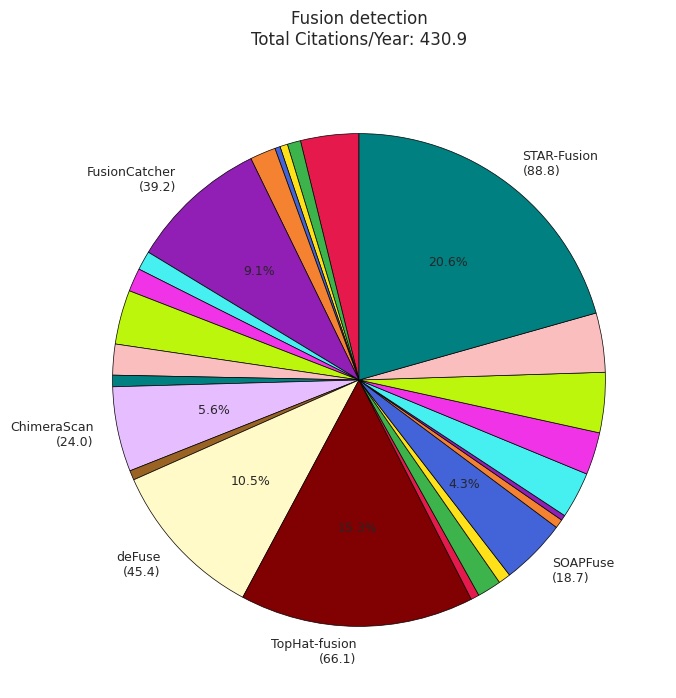

### Figure S1k

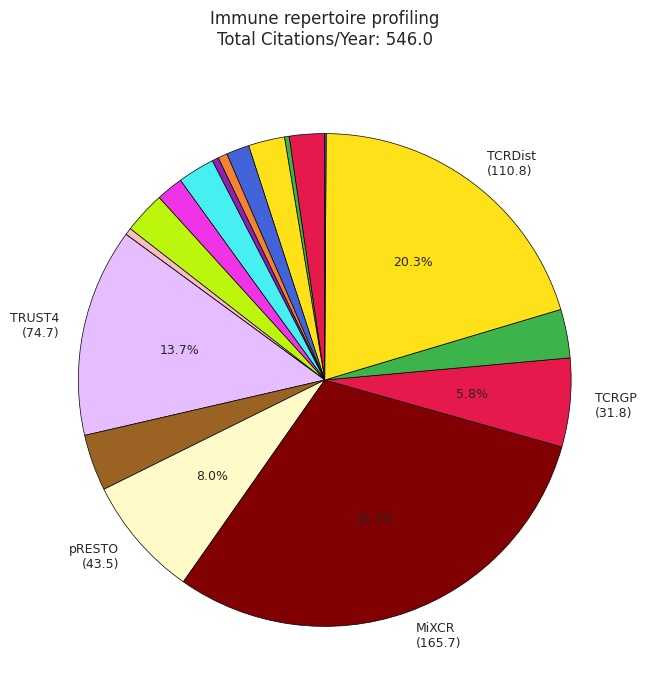

### Figure S1l

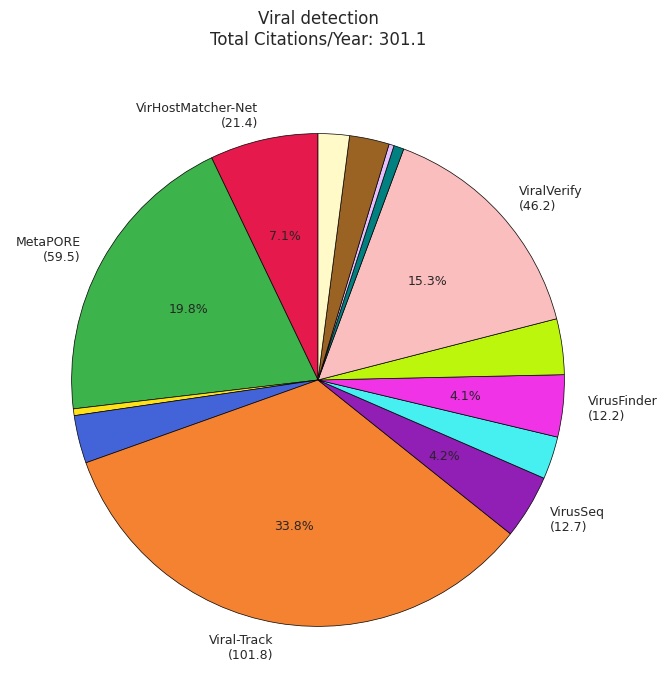

### Figure S1m

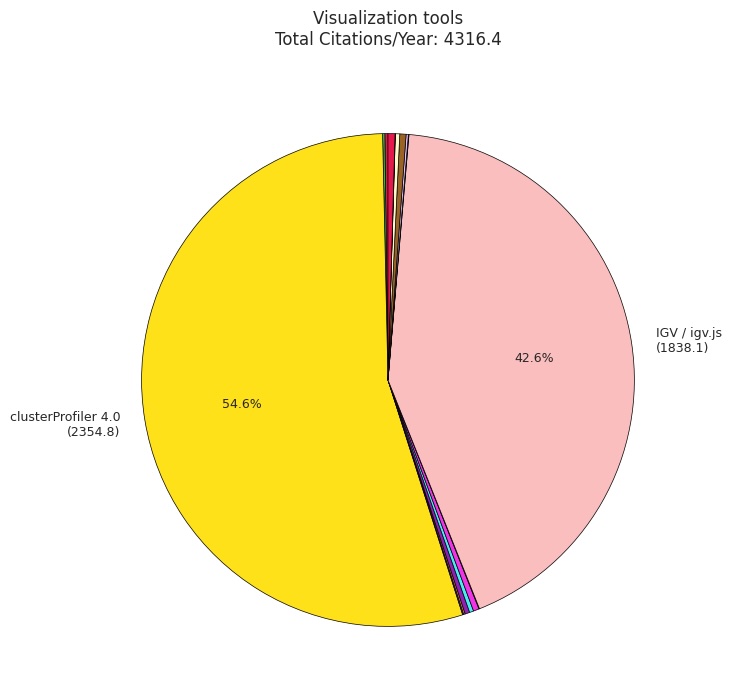

### Figure S1n

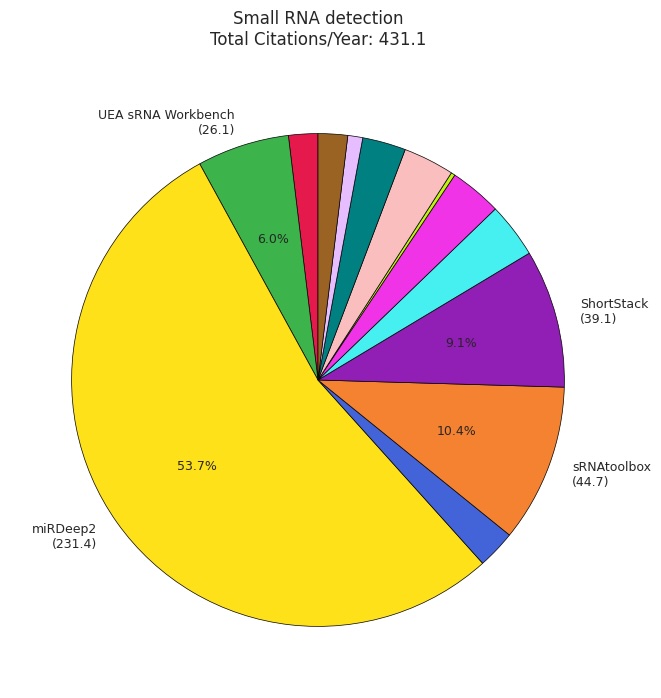

### Figure S1o

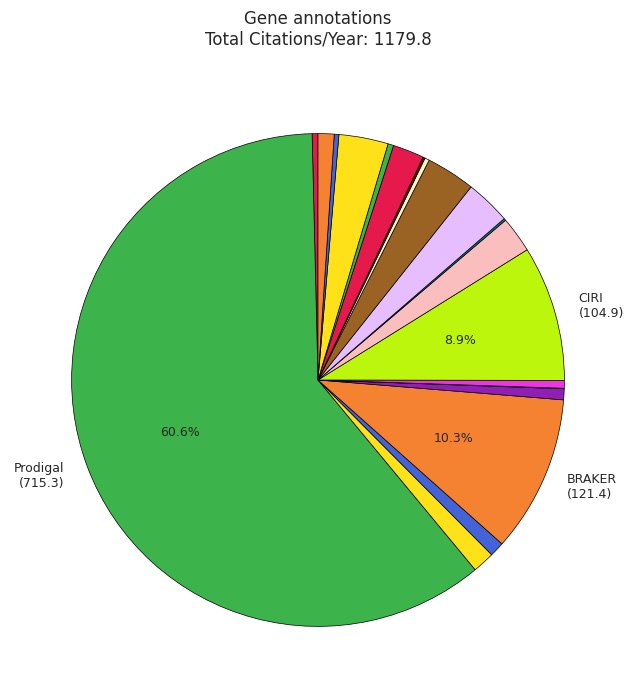

### Figure S2

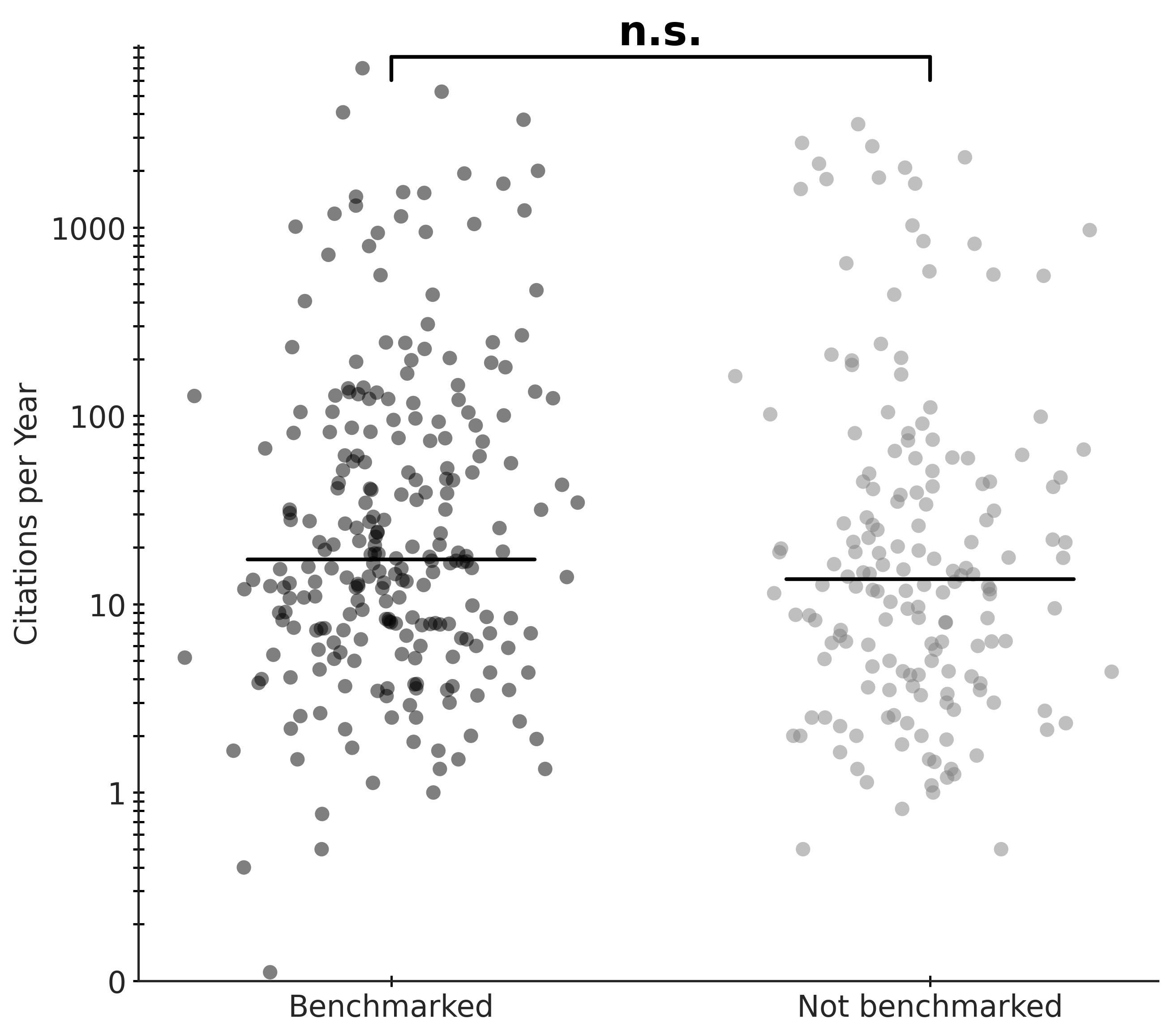

### Figure S3

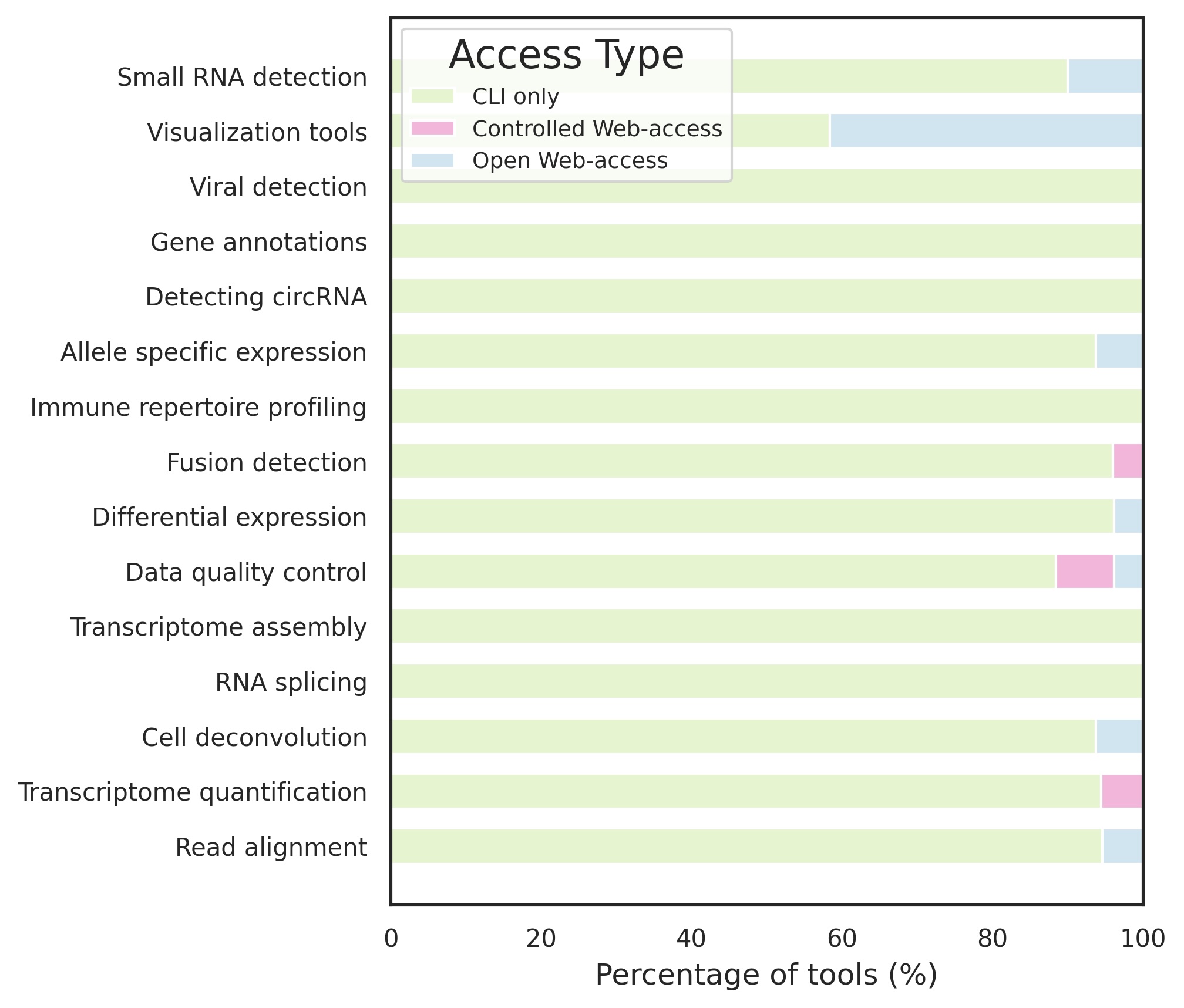

### Figure S4a

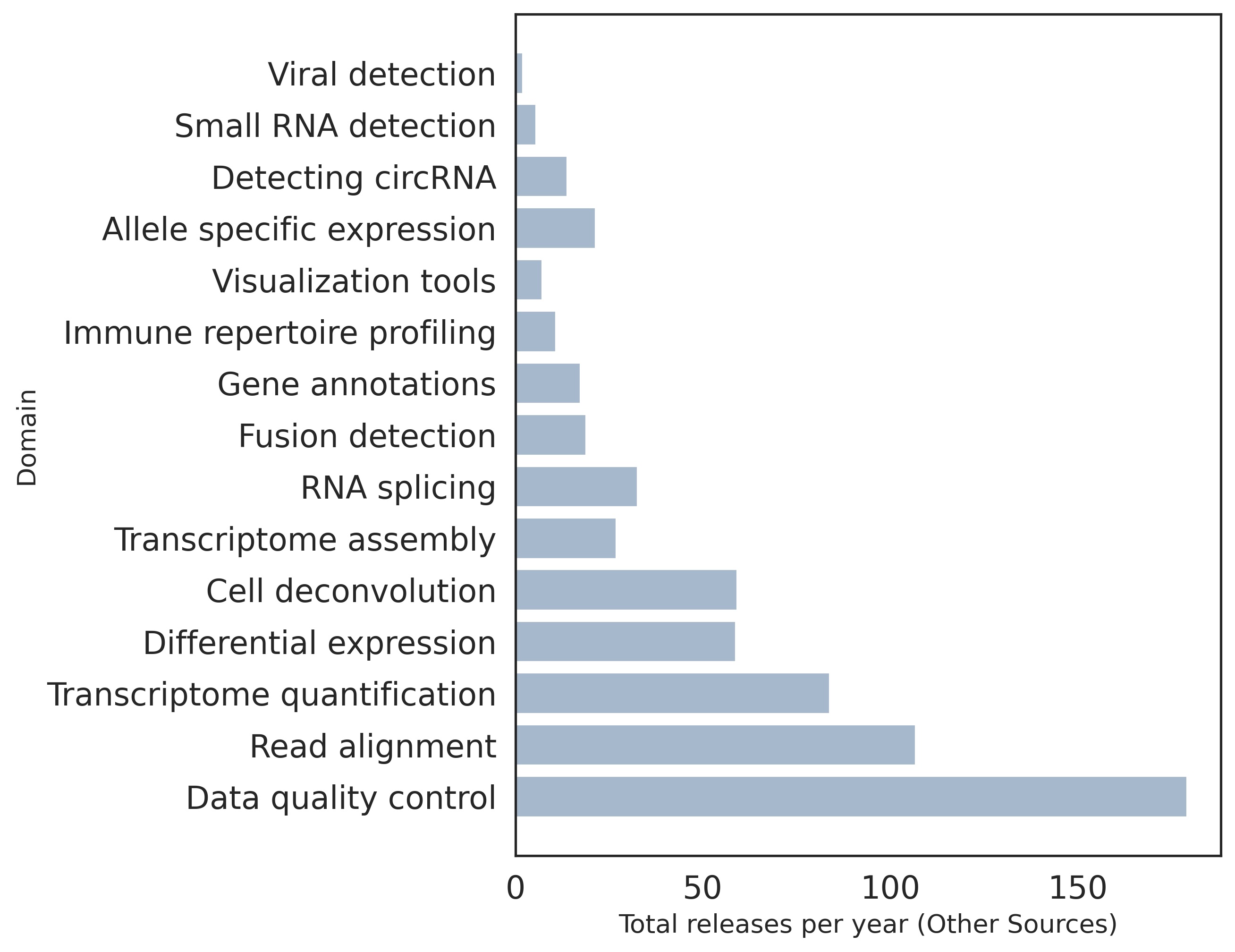

### Figure S4b

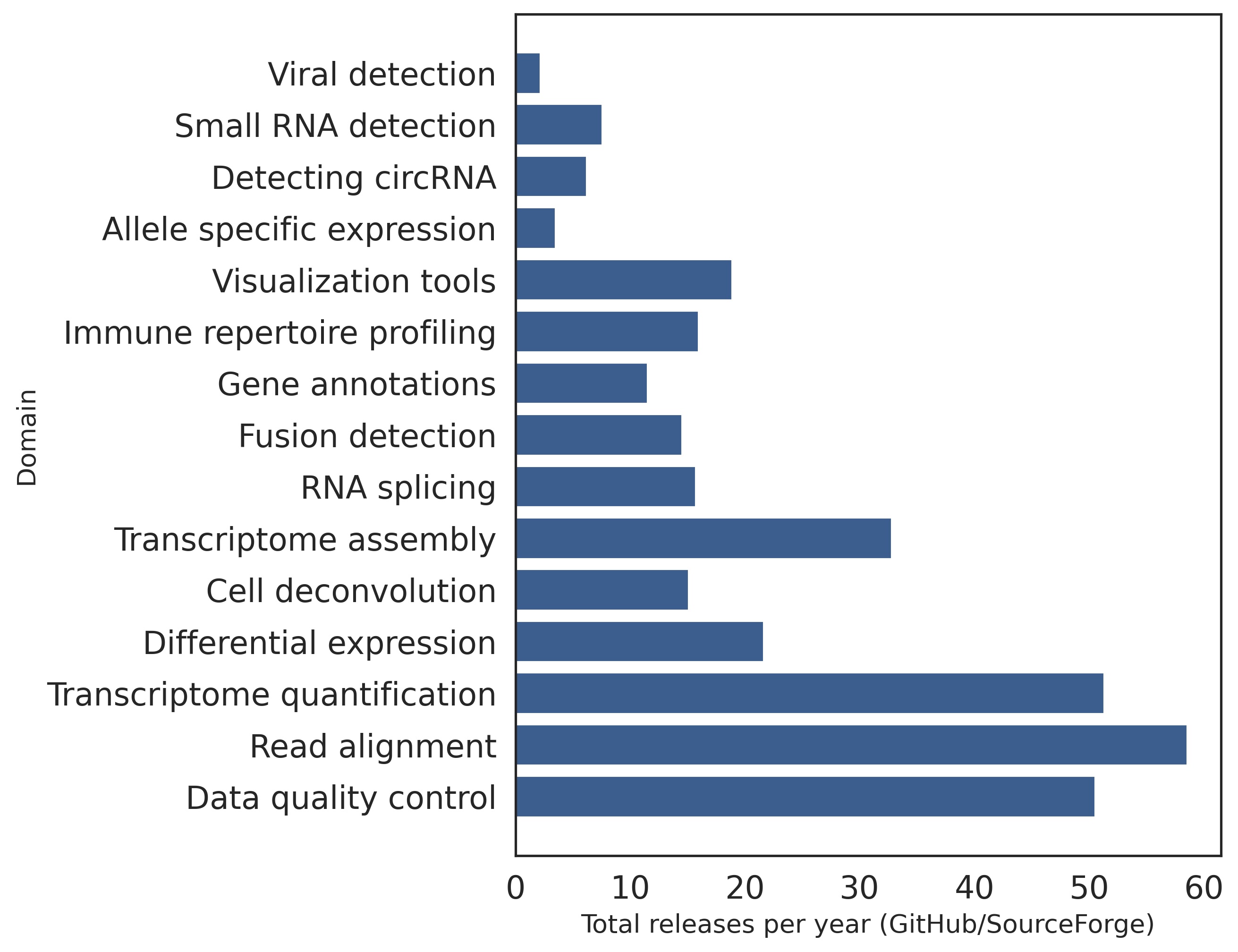

### Figure S6

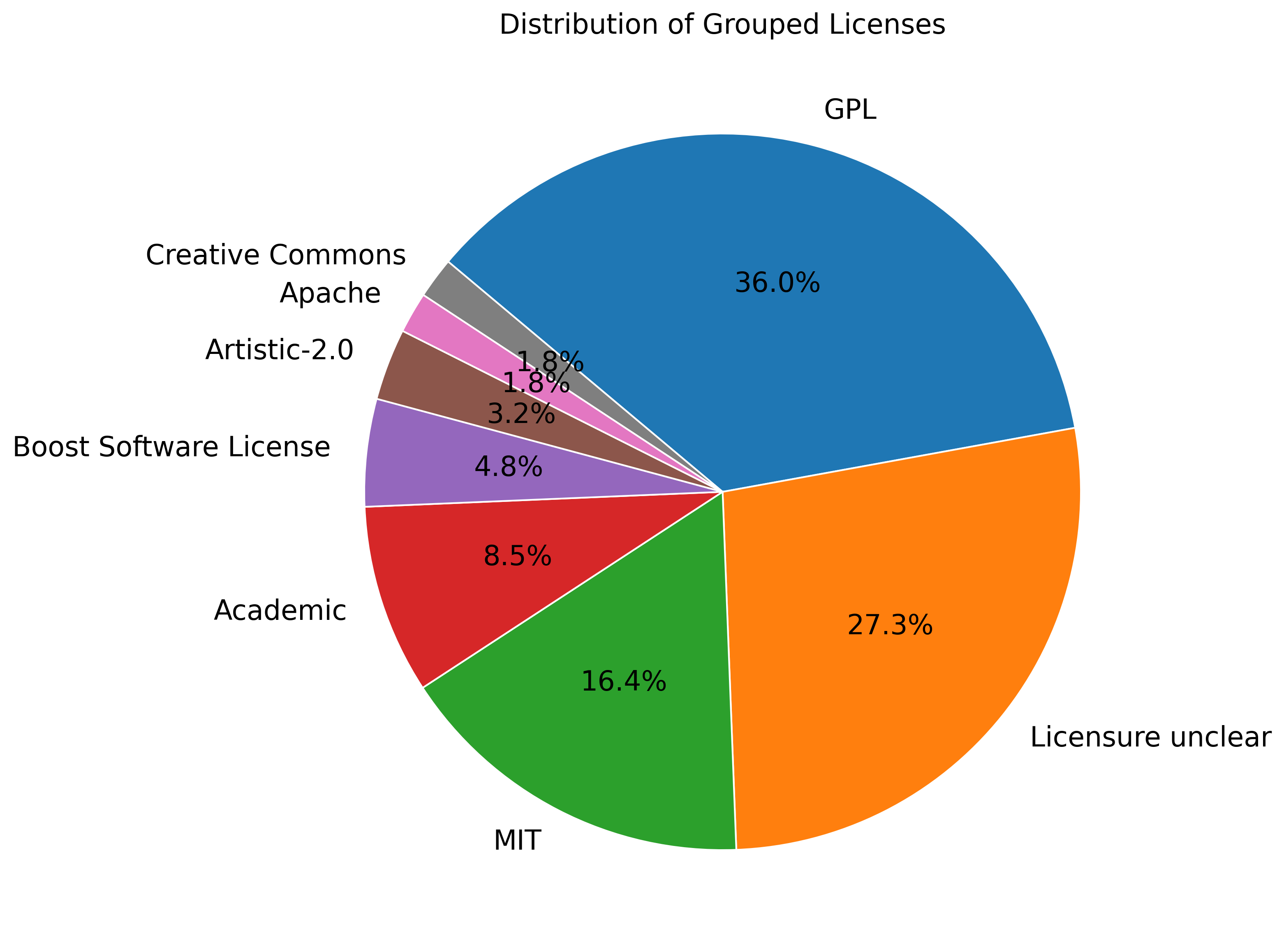

### Figure S7

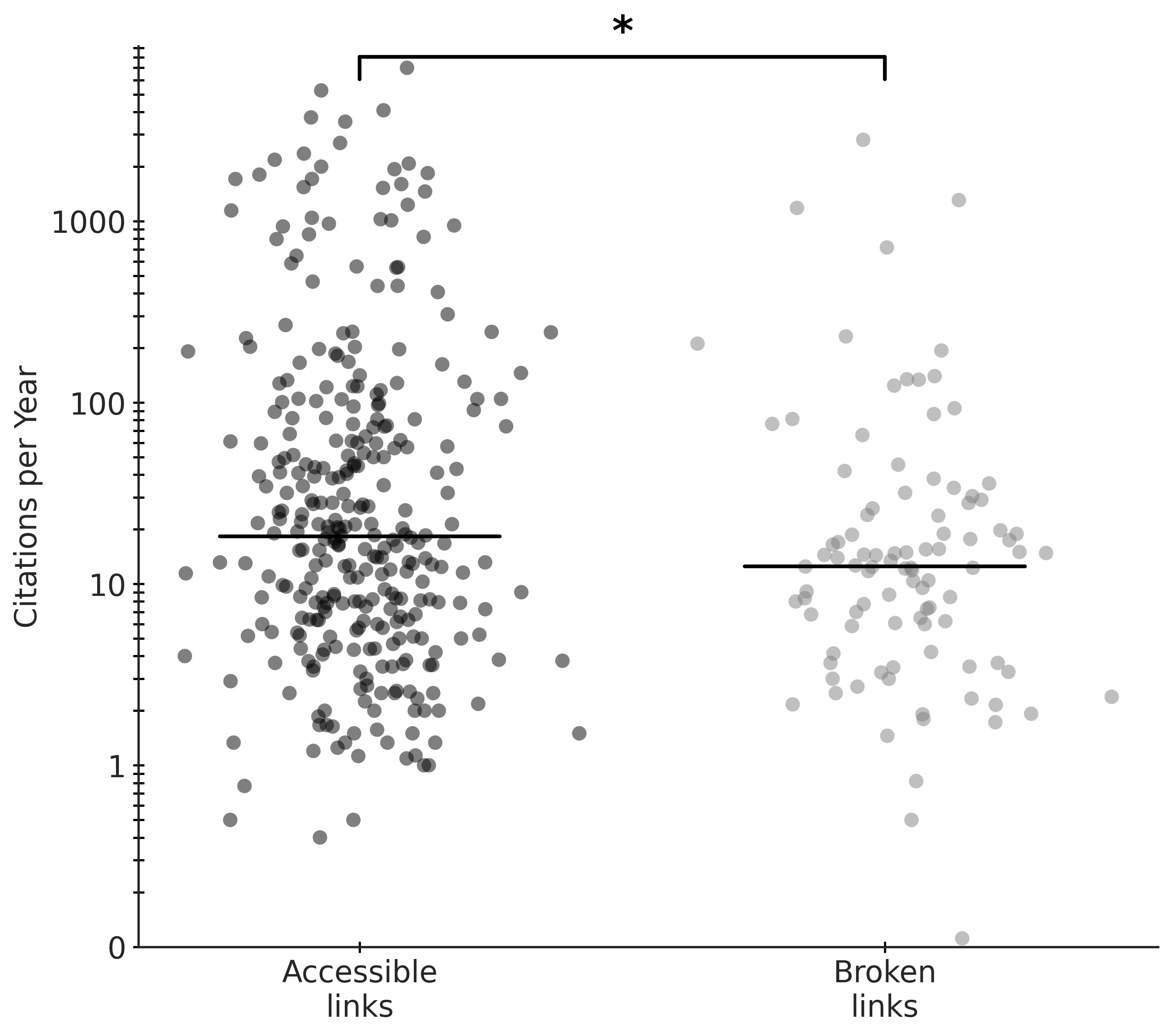

### Figure S8

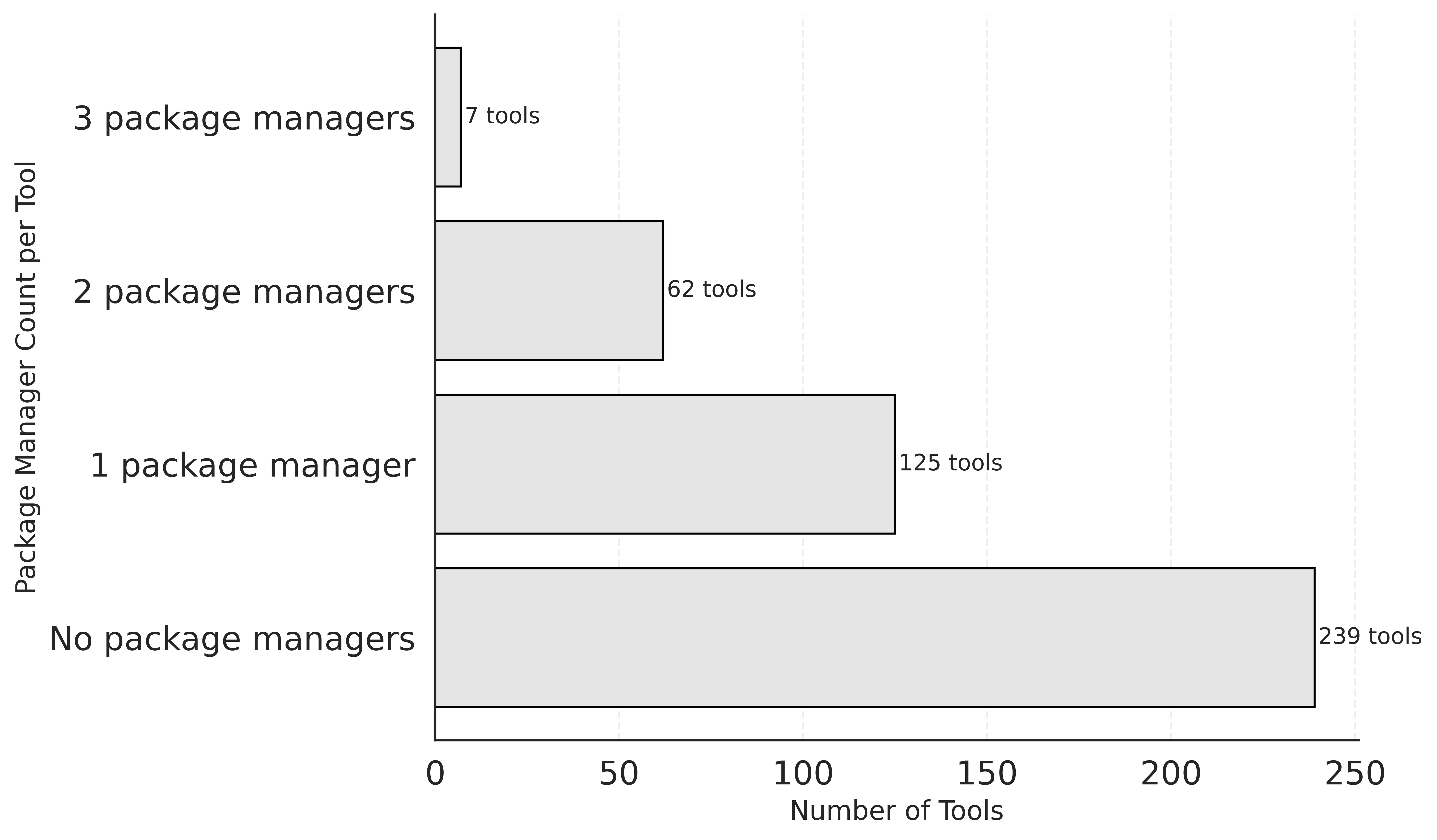

### Figure S9

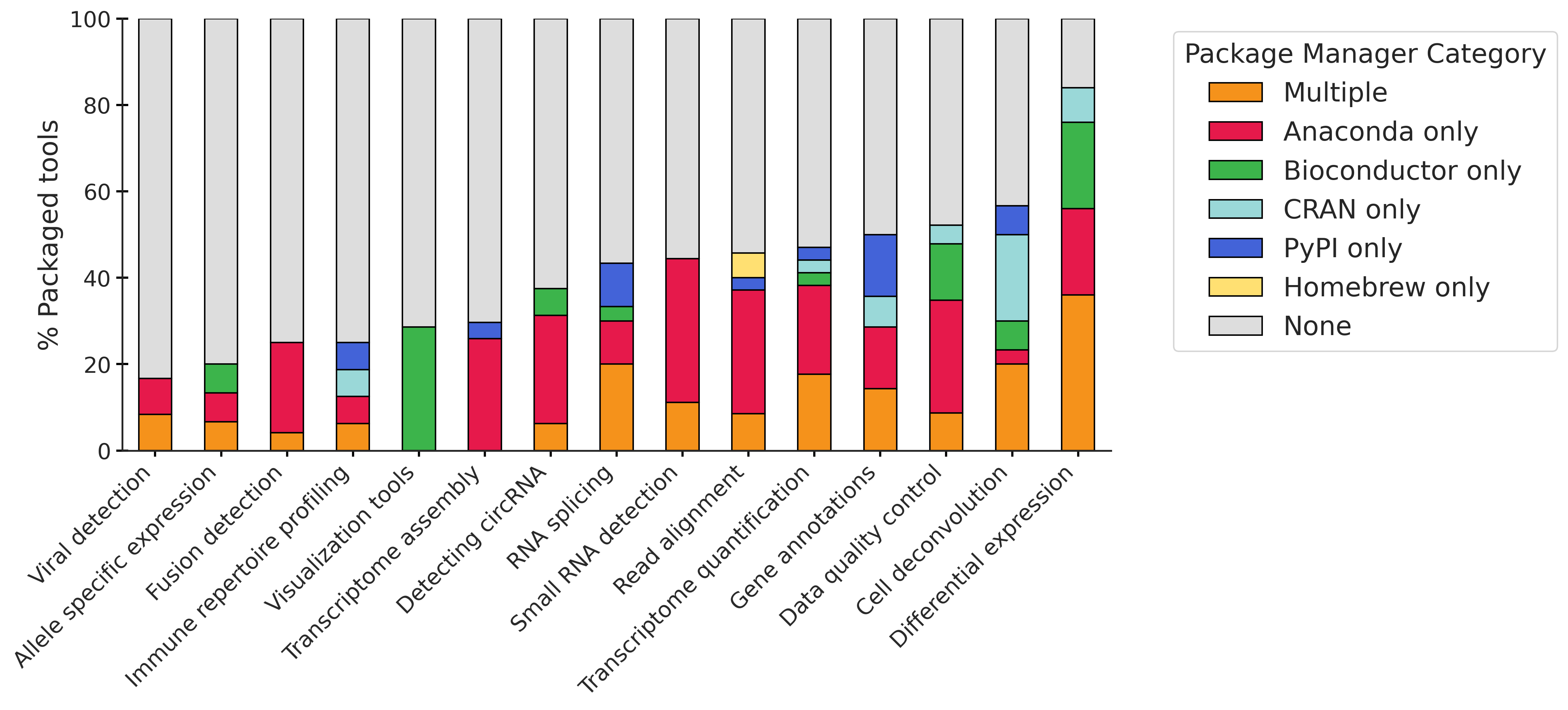

### Figure S10

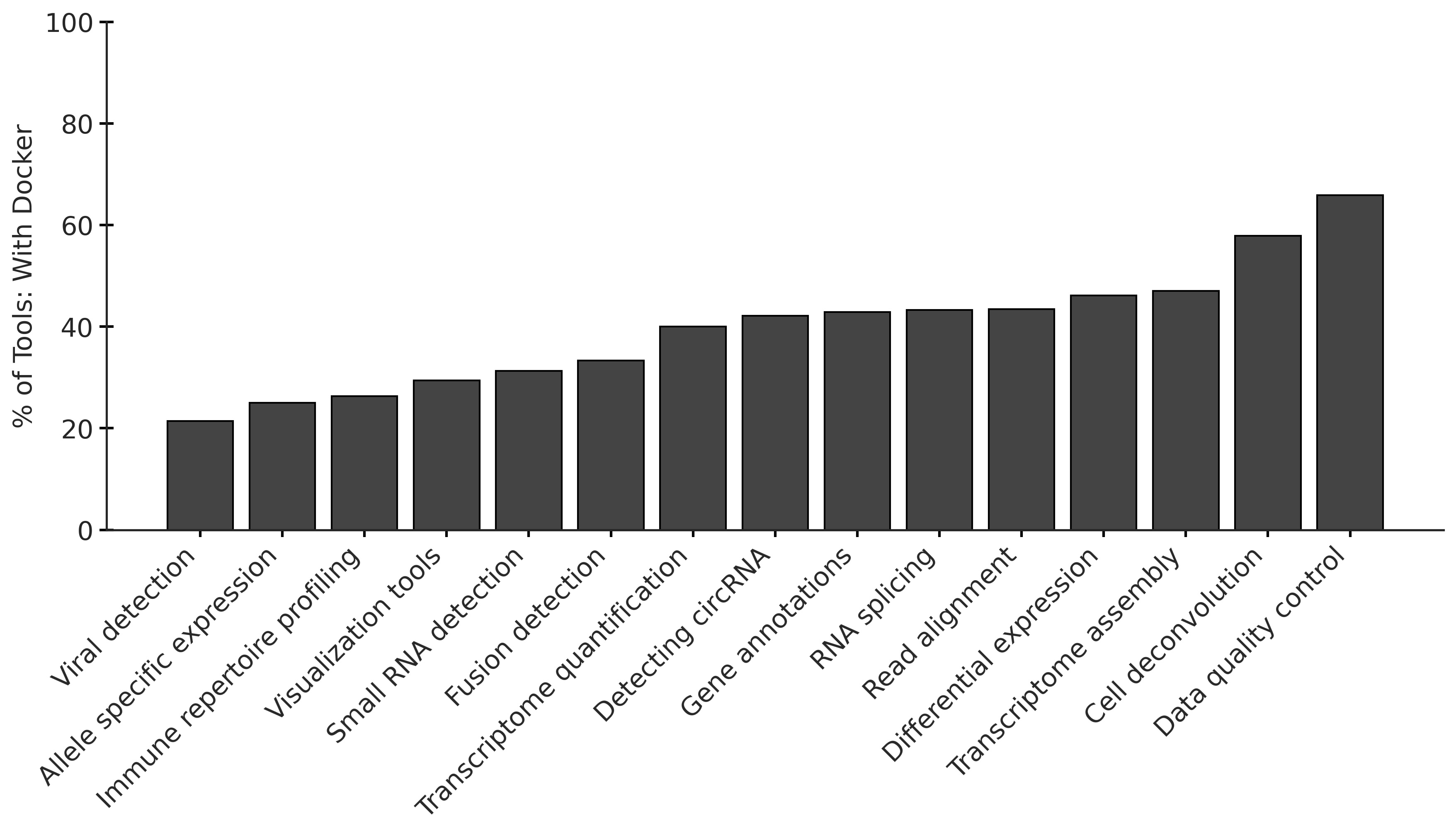
